# The regulation of HAD-like phosphatases by signaling pathways modulates cellular resistance to the metabolic inhibitor, 2-deoxyglucose

**DOI:** 10.1101/504134

**Authors:** Quentin Defenouillère, Agathe Verraes, Clotilde Laussel, Anne Friedrich, Joseph Schacherer, Sébastien Léon

## Abstract

Cancer cells display an altered metabolism with an increased glycolysis and glucose uptake. Anti-cancer strategies targeting glycolysis through metabolic inhibitors have been considered. Particularly, the glucose analogue 2-deoxyglucose (2DG) is imported into cells and phosphorylated into 2DG-6-phosphate, a toxic by-product that inhibits glycolysis. Recent data suggest that 2DG has additional effects in the cell, and resistance to 2DG has also been observed. Using yeast as a model, we engaged an unbiased, mass-spectrometry-based approach to probe the cellular effects of 2DG on the proteome and study resistance mechanisms. This revealed that two 2DG-6-phosphate phosphatases, Dog1 and Dog2, are induced upon exposure to 2DG and participate in 2DG detoxication. 2DG induces Dog2 by upregulating several signaling pathways, such as the MAPK (Hog1/p38)-based stress-responsive pathway, the Unfolded Protein Response (UPR) triggered by 2DG-induced ER stress, and the MAPK (Slt2)-based Cell Wall Integrity pathway. Thus, 2DG-induced interference with cellular signaling rewires the expression of these endogenous phosphatases to promote 2DG resistance. Consequently, loss of the UPR or CWI pathways leads to hypersensitivity to 2DG. In contrast, *DOG2* is transcriptionally repressed by glucose availability in a Snf1/AMPK-dependent manner, and mutants impaired in this pathway are 2DG-resistant. The characterization and genome resequencing of spontaneous 2DG-resistant mutants revealed that *DOG2* overexpression is a common strategy to achieve 2DG resistance. The human Dog2 orthologue, HDHD1, also displays 2DG-6-phosphate phosphatase activity *in vitro*, and its overexpression confers 2DG resistance in HeLa cells, which has important implications for potential future chemotherapies involving 2DG.

## Introduction

Most cancer cells display an altered metabolism, with an increased glucose consumption to support their proliferative metabolism that is based on aerobic glycolysis (Warburg effect) (*1, 2*). Targeting glycolysis has been proposed as a strategy to target cancer cells and various metabolic inhibitors have been considered (*3, 4*).

2-deoxy-D-glucose (2DG) is a derivative of D-glucose that is actively imported by glucose transporters and is phosphorylated by hexokinase into 2-deoxy-D-glucose-6-phosphate (2DG6P), but cannot be further metabolized due to the 2-deoxy substitution, triggering a decrease in cellular ATP content in tumors (*5*). Mechanistically, 2DG6P accumulation hampers glycolysis by inhibiting hexokinase activity in a non-competitive manner (*6, 7*), as well as phospho-glucose isomerase activity in a competitive manner (*8*). Since cancer cells rely on an increased glycolysis rate for proliferation, 2DG has been of interest for cancer therapy, particularly in combination with radiotherapy or other metabolic inhibitors (*9–11*). These features led to a phase I clinical trial using 2DG in combination with other drugs to treat solid tumors (*12*). Its derivative ^18^Fluoro-2DG is also used in cancer imaging (PET scans) as it preferentially accumulates in tumor cells due to their increased glucose uptake (*13*). Additionally, due to its structural similarity to mannose, 2DG (which could also be referred as to 2-deoxymannose, since mannose is the C2 epimer of glucose) also interferes with N-linked glycosylation and causes ER stress (*14–16*) and this was proposed to be the main mechanism by which 2DG kills normoxic cells (*17*). Recently, 2DG toxicity was also linked to the depletion of phosphate pools following 2DG phosphorylation (*18*). Finally, interference of 2DG with lipid metabolism and calcium homeostasis was also described, but the underlying mechanisms are less clear (*19*). Intriguingly, despite its pleiotropic mode of action, resistance to 2-deoxyglucose was reported in cell cultures (*20*).

Because these metabolic and signalling pathways are evolutionarily conserved, simpler eukaryotic models such as the budding yeast *Saccharomyces cerevisiae* can be used to understand the mode of action of 2DG. Moreover, yeast is particularly well-suited for these studies because of its Warburg-like metabolism (*21*). Akin to cancer cells, *Saccharomyces cerevisiae* privileges glucose consumption through glycolysis over respiration, regardless of the presence of oxygen. This is allowed by a glucose-mediated repression of genes involved in respiration and alternative carbon metabolism, which operates at the transcriptional level. This glucose-mediated repression mechanism is relieved upon activation of the yeast orthologue of AMPK, Snf1, which phosphorylates the transcriptional repressor Mig1 and leads to its translocation out of the nucleus (*22–25*). Previous studies in yeast identified mutations that render yeast cells more tolerant to 2DG (*26–31*). This suggested the existence of cellular mechanisms that can modulate 2DG toxicity, which are important to characterize if 2DG were to be used for therapies.

In yeast, 2DG was initially used to identify genes involved in glucose repression. This was based on the observation that 2DG, like glucose, causes Snf1 inactivation and thus prevents the use of alternative carbon sources (*32–34*). The characterization of mutants that were able to grow in sucrose medium despite the presence of 2DG allowed the identification of actors of the glucose repression pathway (*26, 29, 35*). These experiments also revealed that mutations in *HXK2*, encoding hexokinase II, also rendered yeast cells more tolerant to 2DG, perhaps by limiting 2DG phosphorylation and 2DG6P accumulation (*29–31, 36, 37*). Finally, several 2DG-resistant mutants displayed an increased 2DG6P phosphatase activity, which could detoxify the cells of this metabolite and dampen its negative effects on cellular physiology (*38, 39*). Indeed, two 2DG6P phosphatases were subsequently cloned, named *DOG1* and *DOG2*, and their overexpression led to 2DG resistance and prevented the 2DG-mediated repression of genes (*33, 40, 41*).

More recently, the toxicity of 2DG was studied in the context of glucose-grown cells, which may be more relevant for the understanding its mode of action in mammalian cells. In these conditions, 2DG toxicity is independent of its effect on the glucose repression of genes, but involves distinct mechanisms, such as a direct inhibition of glycolysis and other cellular pathways (*30, 42, 43*). Accordingly, several mutations initially identified as leading to 2DG tolerance in sucrose medium have no effect in glucose medium (*30, 44*). A key finding was that the deletion of *REG1*, encoding a regulatory subunit of Protein Phosphatase 1 (PP1) that negatively regulates Snf1 (*45, 46*) leads to 2DG resistance (*30*). The resistance of the *reg1∆* mutant depends on the presence of Snf1, and the single deletion of *SNF1* also renders yeast hypersensitive to 2DG (*31*). These data demonstrate that Snf1 activity is crucial for 2DG resistance. A model was proposed in which the 2DG sensitivity displayed by the *snf1∆* mutant involves a misregulated expression and localization of the low-affinity glucose transporters, Hxt1 and Hxt3 (*47*). Additionally, the deletion of *LSM6*, which encodes a component of a complex involved in mRNA degradation, also leads to 2DG resistance in a Snf1-dependent manner, but the mechanism by which this occurs is unknown (*31*). Thus, many aspects of the pathways mediating 2DG sensitivity/resistance remain to be explored.

In the present study, we engaged an unbiased, mass-spectrometry-based approach in yeast to better understand the cellular response to 2DG. This revealed that the main 2DG6P phosphatase, Dog2, is induced upon exposure to 2DG and participates in 2DG detoxification in glucose medium. We reveal that 2DG induces Dog2 (and Dog1, to a certain extent), by upregulating several stress-responsive signaling pathways (UPR and MAPK-based pathways). Moreover, the expression of *DOG2* is additionally regulated by Snf1 and the glucose-repression pathway through the action of downstream transcriptional repressors, and contributes to the resistance of glucose-repression mutants to 2DG. The partial characterization of 24 spontaneous 2DG-resistant mutants revealed that most display an increased *DOG2* expression, suggesting it is a common strategy used to acquire 2DG resistance. Particularly, genome resequencing identified that mutations in *CYC8*, encoding a transcriptional corepressor, cause 2DG resistance through the upregulation of Dog2. The identification of a potential human orthologue of the Dog1/2 proteins, named HDHD1, which displays an *in vitro* 2-DG-6-phosphate phosphatase activity and which can cause resistance of HeLa cells to 2DG when overexpressed, suggests that HAD-like phosphatases are conserved regulators of 2DG resistance.

## Results

### A proteomic assessment of the cellular response to 2DG reveals an increased expression of metabolic enzymes

As a first step to characterize the cellular response of yeast cells to 2-deoxyglucose (2DG) treatment in an unbiased and quantitative manner, we performed proteome-wide mass spectrometry-based proteomics on wild-type cells treated for 2.5 hours with 0.2% 2DG, a concentration that prevents growth of WT cells on plates (*30*) using untreated cells as a negative control. Overall, 78 proteins were significantly up-regulated, whereas 18 proteins were down-regulated after 2DG treatment (False Discovery Rate of 0.01) (Figure 1A, **Table S1**). Among the up-regulated candidates, a significant enrichment of proteins involved in various metabolic processes was noted, including that of glucose, glucose-6-phosphate and other carbohydrates (Figure 1B, Figure S1A) in line with the fact that 2DG interferes with glycolysis (*5*).

**Figure 1.**
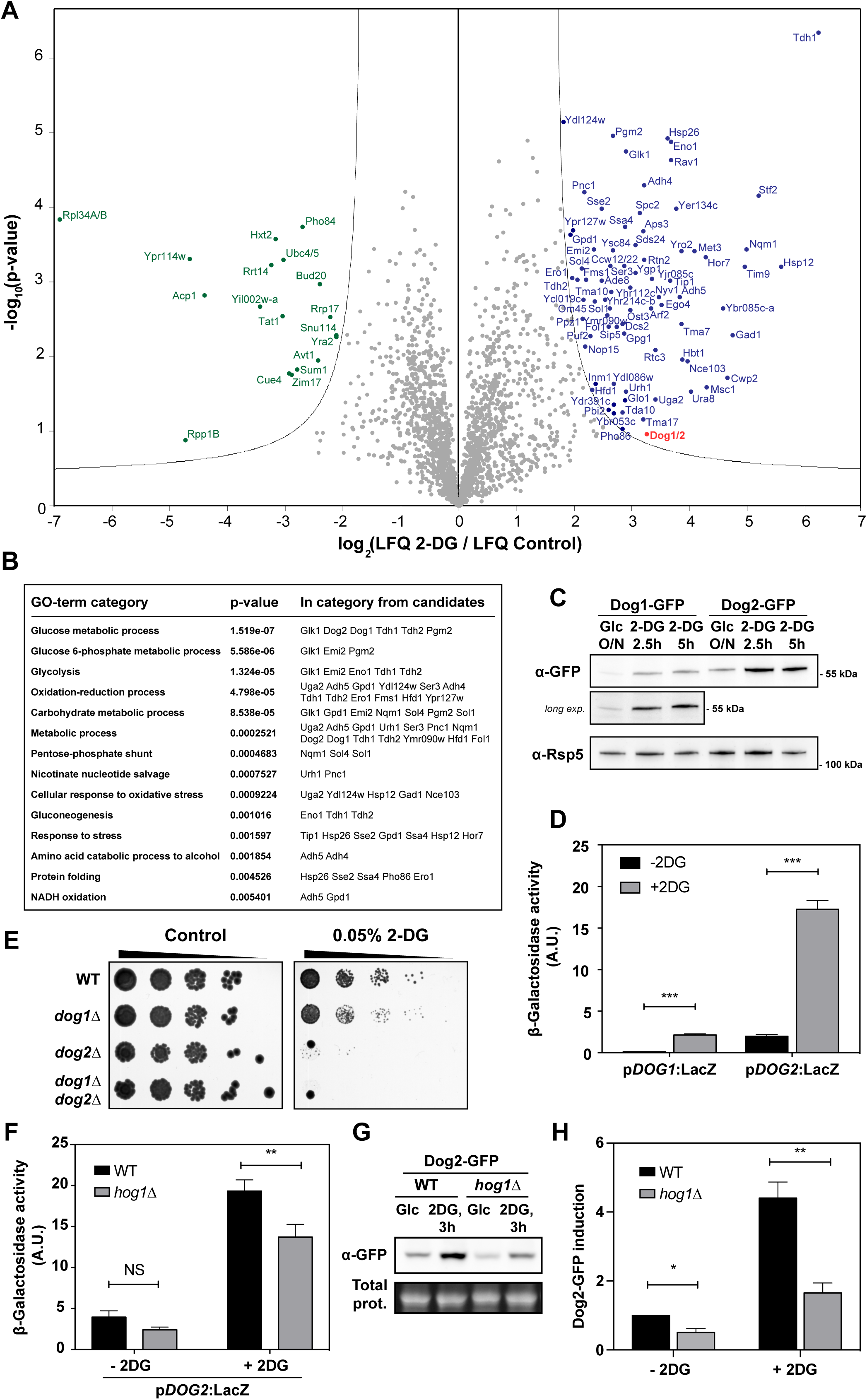
Proteomics analysis of yeast response to 2DG reveals a transcriptional induction of the 2DG6P phosphatases Dog1-Dog2. (**A**) Volcano-plot representing changes in protein abundance in total protein extracts of wild-type (WT) yeast in response to 2-deoxyglucose (0.2%), obtained by mass spectrometry-based proteomics and analyzed with the MaxQuant software. The x axis corresponds to the log_2_ value of the abundance ratio (LFQ: Label-Free Quantification) between 2DG treatment and the negative control. The y axis represents the −log_10_ of the *p*-value of the statistical t-test for each quantified protein (n=3 independent biological replicates). Line: threshold with a False Discovery Rate of 0.01. (**B**) Gene Ontology (GO) analysis of the proteins identified as upregulated in response to 2DG treatment along with their *p-*value and the proteins included in each category. (**C**) Western blot on total protein extracts of yeast cells expressing endogenously tagged Dog1-GFP or Dog2-GFP, before and after 2DG addition for the indicated time, using an anti-GFP antibody. A longer exposure is displayed for Dog1-GFP cells to highlight the higher abundance of Dog1 after 2DG addition. Rsp5, whose levels did not change upon 2DG addition in all of our experiments, is used as a loading control. (**D**) Beta-galactosidase assays of wild-type yeast cells expressing *LacZ* under the control of the p*DOG1* or p*DOG2* promoters, before and after 2DG treatments for 3 hours (± SEM, *n*=3). (**E**) Serial dilutions of cultures from the indicated strains were spotted onto SD plates containing no DG or 0.05% 2DG, and grown for 3 days at 30°C. (**E**) Beta-galactosidase assays of wild-type and *hog1∆* strains expressing *LacZ* under the control of the p*DOG2* promoter, before and after 2DG treatments for 3 hours. Error bars: SEM (*n*=3). (**F**) Western blot on total protein extracts of WT and *hog1∆* cells endogenously expressing a Dog2-GFP fusion, before and after 2DG addition for 3h, using an anti-GFP antibody. Total proteins were visualized in gels using a trihalo compound (see *Methods*). (**G**) Relative expression of Dog2-GFP in the same conditions as (**F**) after normalization by total proteins and using WT/untreated as a reference (± SEM, *n*=3).

Interestingly, these proteomics data revealed an increase in the abundance of the 2-deoxyglucose-6-phosphate phosphatases Dog1 or Dog2, which could not be discriminated at the mass spectrometry level because of their high sequence identity (92%). Although the expression of these phosphatases was previously shown to promote 2DG resistance (*33, 40, 41*), the increased expression of Dog1 and/or Dog2 in response to 2DG was intriguing because it raised the question of how exposure to this synthetic molecule could trigger an adaptive resistance mechanism in yeast. To first confirm that 2DG induces Dog1 and Dog2, they were individually GFP-tagged at their chromosomal locus, in order to maintain an endogenous regulation, and their expression was monitored by western blotting. This revealed that both Dog1 and Dog2 expression levels are increased in the presence of 2DG, and that Dog2-GFP is more abundant than Dog1-GFP (Figure 1C).

To evaluate the contribution of transcription in the regulation of the *DOG1* and *DOG2* genes by 2DG, the corresponding promoters (1kb) were fused to a beta-galactosidase reporter. This revealed an increased activity of the *DOG1* and *DOG2* promoters in response to 2DG (Figure 1D), suggesting that the expression of Dog1 and Dog2 is, at least in part, due to an increased transcription. Of note, the deletion of the *DOG2* gene strongly sensitized yeast cells to 2-deoxyglucose, but that of *DOG1* had little effect (Figure 1E), indicating that Dog2 is functionally more important than Dog1, perhaps due to its higher expression level (Figure 1C). This indicates that Dog2 participates to the natural tolerance of WT cells to low concentrations (0.05%) of 2DG.

Prior work showed that the expression of Dog2, whose endogenous function in yeast metabolism is unknown, is induced by various stresses, such as oxidative and osmotic stresses, through the stress-responsive MAP kinase orthologue of p38, Hog1 (*48*). The deletion of *HOG1* prevented the maximal induction of *pDOG2*-LacZ expression (Figure 1F) and Dog2 induction (Figure 1G-H), but had no discernable effect on the expression of Dog1 (Figure S2A). When we tested the effect of 2DG on Hog1 phosphorylation using an antibody directed against the phosphorylated form of mammalian p38 (*49*), we found that Hog1 appeared partially activated upon 2DG addition, but to a much lower extent as observed after hyperosmotic shock (Figure S2B). These data show that the stress-activated protein kinase Hog1 participates in Dog2 induction but also that the maximal upregulation of Dog1 and Dog2 by 2DG involves at least one additional level of regulation.

### The expression of Dog1 and Dog2 is induced by the Unfolded Protein Response pathway through 2-deoxyglucose-induced ER stress

Exposure of cancer cells to 2-deoxyglucose interferes with N-linked glycosylation, likely because of the structural similarity of 2-deoxyglucose with mannose, a constituent of the N-glycan structures (*14*). This results in endoplasmic reticulum (ER) stress and consequently, in the induction of the Unfolded Protein response (UPR) pathway in mammalian cells (*14*). Accordingly, the glucose fluorescent analogue 2-NBDG ((2-(*N*-(7-Nitrobenz-2-oxa-1,3-diazol-4-yl)Amino)-2-Deoxyglucose), which was previously used in yeast as a glucose tracer (*50*) and which, much like 2DG, cannot be fully metabolized because of the lack of a hydroxyl group in position 2, partially co-localized with an ER marker (Figure 2A), suggesting interference with N-glycosylation which takes place at this organelle. Treatment of yeast with 2DG also induced a defect in the glycosylation of carboxypeptidase Y (CPY), a vacuolar (lysosomal) protease whose membrane-anchored precursor is N-glycosylated in the course of its intracellular trafficking (*51*). This defect was not as extensive as that observed upon treatment of cells with tunicamycin, an inhibitor of the first step of glycosylation that also causes ER stress and is a strong UPR inducer (*52, 53*) (Figure 2B). Interestingly, the addition of exogenous mannose in the medium suppressed the glycosylation defects caused by 2DG, but not those caused by tunicamycin (Figure 2B). This supports the idea that 2DG mimics mannose and interferes with its incorporation in N-glycans, as proposed in mammalian cells (*14, 17*).

**Figure 2.**
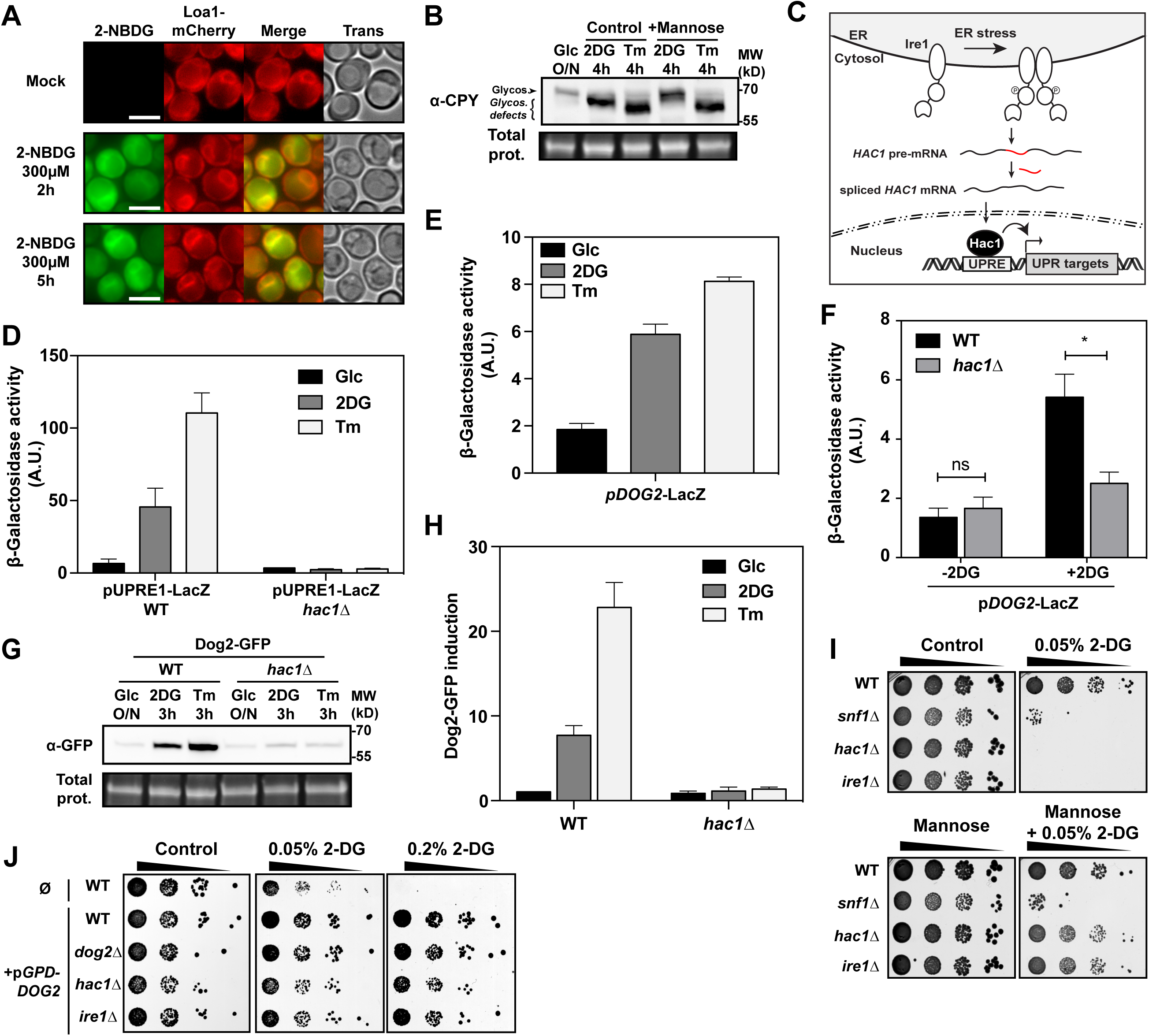
2DG treatment induces Dog2 expression through glycosylation defects that trigger ER stress and the Unfolded Protein Response. (**A**) Cells expressing endogenously-tagged Loa1-mCherry, used as a control of ER localization (*115*), were grown overnight in SC medium (exponential phase) and treated with 300µM 2NBDG (see *Methods*) for the indicated times and imaged by fluorescence microscopy. Scale bar: 5 µm. (**B**) WT cells were grown overnight to mid-log phase in SC medium, centrifuged and resuspended in SC-medium containing mannose (2%) or not, and treated with 0.2% 2DG or 1µg/mL tunicamycin for 4 hours. Total protein extracts were prepared and a western blot on was done using anti-Carboxypeptidase Y (CPY) antibodies. (**C**) Beta-galactosidase assays on WT and *hac1∆* cells expressing *LacZ* under the control of an UPR-inducible promoter (p*UPRE1*) and treated with 0.2% 2DG or 1µg/mL tunicamycin for 3 hours (± SEM, *n*=3). (**D**) Beta-galactosidase assays on WT cells expressing *LacZ* under the control of the *DOG2* promoter and treated as in (**C**) (± SEM, *n*=3). (**E**) Beta-galactosidase assays on WT and *hac1∆* cells expressing *LacZ* under the control the *DOG2* promoter, before and after 3h 2DG treatments (± SEM, *n*=3). (**F**) Western blot on total protein extracts of WT and *hac1∆* cells endogenously expressing a Dog2-GFP fusion, before and after 3h treatment with 2DG or tunicamycin, using an anti-GFP antibody. (**G**) Relative expression of Dog2-GFP in the same conditions as (**F**) after normalization by total proteins and using WT/untreated as a reference (± SEM, *n*=3). (**H**) Serial dilutions of cultures from the indicated strains were spotted onto SC plates (supplemented with 2% mannose when indicated) containing no DG or 0.05% 2DG, and were grown for 3 days at 30°C. (**I**) Serial dilutions of cultures from the indicated strains overexpressing *DOG2* (*pGPD*-*DOG2*) or not (Ø) were spotted onto SC-Ura plates containing 0, 0.05% or 0.2% 2-deoxyglucose. The plates were scanned after 3 days of incubation at 30°C.

In yeast, the UPR pathway is initiated by a multifunctional ER membrane protein named Ire1. Upon sensing ER stress and after dimerization, Ire1 will splice pre-mRNAs encoding the Hac1 transcription factor, leading to its translation and the trans-activation of Hac1 targets carrying an UPRE (UPR Response Element) in their promoter (Figure 2C) (*54*). To address whether 2DG can induce the UPR pathway in yeast, we measured the induction of an UPRE-driven reporter, UPRE1-LacZ (*53*) in response to 2DG treatment. Not only was 2DG able to elicit the expression of this reporter, but this depended on Hac1, confirming that 2DG is a *bona fide* UPR inducer (Figure 2D).

We then tested whether 2DG-mediated induction of Dog2 involves the UPR pathway. First, we found that tunicamycin strongly induced the *pDOG2*-LacZ reporter (Figure 2E), suggesting that glycosylation defects and/or the ensuing ER stress promotes *DOG2* expression. Moreover, the induction of this reporter by 2DG was reduced by more than 2-fold in a *hac1∆* mutant, suggesting a contribution of the UPR in this induction (Figure 2F). Comparable results were obtained for p*DOG1*-LacZ (Figure S3). These results were confirmed at the protein level for Dog2 (Figure 2G-H).

Interestingly, UPR-compromised mutants such as *hac1∆* and *ire1∆* are both hypersensitive to 2DG (Figure 2I), suggesting that they cannot cope with ER stress caused by 2DG. The addition of exogenous mannose in the medium, that we showed can alleviate the glycosylation defects caused by 2DG (see Figure 2B), restored 2DG-tolerance of these mutants to a comparable level as the WT (Figure 2I) but not that of *snf1∆*, a known 2DG-hypersensitive mutant (*31*) (see also Figure 4D). Altogether, these results confirm that 2DG induces ER stress by interfering with N-glycosylation, that the subsequent activation of the UPR pathway upregulates Dog1 and Dog2 expression and that UPR mutants display a lower level of Dog1 and Dog2 expression. Noticeably, Dog2 overexpression restored the growth of *hac1∆* or *ire1∆* cells at various concentrations of 2DG (Figure 2J).

### 2DG also activates the MAPK-based cell-wall integrity pathway which additionally contributes to Dog1 and Dog2 expression

To address a possible additional contribution of other signaling pathways that would contribute to the 2-DG-induced expression of Dog1 and Dog2, we ran a bioinformatics analysis on the list of proteins that were found as upregulated upon 2DG treatment (see Figure 1A, and **Table S1**) using YEASTRACT (*55*) to evaluate whether the observed variations in protein expression could reveal a potential transcriptional signature. The best hit (corresponding to 35/78 candidates, *p-*value=0) was the MADS-box transcription factor Rlm1, a known downstream target of the cell wall integrity (CWI) signaling pathway (*56*). The cell wall integrity pathway is activated by several stresses such as cell wall damage, and involves plasma membrane-localized sensors, the Rho1 GTPase and its cofactors (Guanine nucleotide exchange Factors [GEFs] and accessory proteins), protein kinase C, and a cascade leading to the activation of the MAPK Slt2 and downstream transcription factors, such as Rlm1 (Figure 3A) (*56*). There are well-established links between CWI signaling and ER stress (*57–61*); in particular, the CWI pathway was found to be required for viability when facing tunicamycin-induced ER stress. Importantly, we found that the deletion of many genes of the CWI pathway acting upstream of Rlm1 caused an increased 2DG sensitivity (Figure 3B) such as those encoding the sensors *MID2* and *WSC1*. Thus, much like the UPR pathway, the CWI pathway is required for 2DG tolerance. Importantly, 2DG treatment induced the phosphorylation of the MAPK Slt2 within the first hour of exposure (Figure S4A). 2DG also triggered the induction of an Rlm1-regulated promoter fused to LacZ (Figure S4B). Thus, the CWI pathway is activated by 2DG treatment.

**Figure 3.**
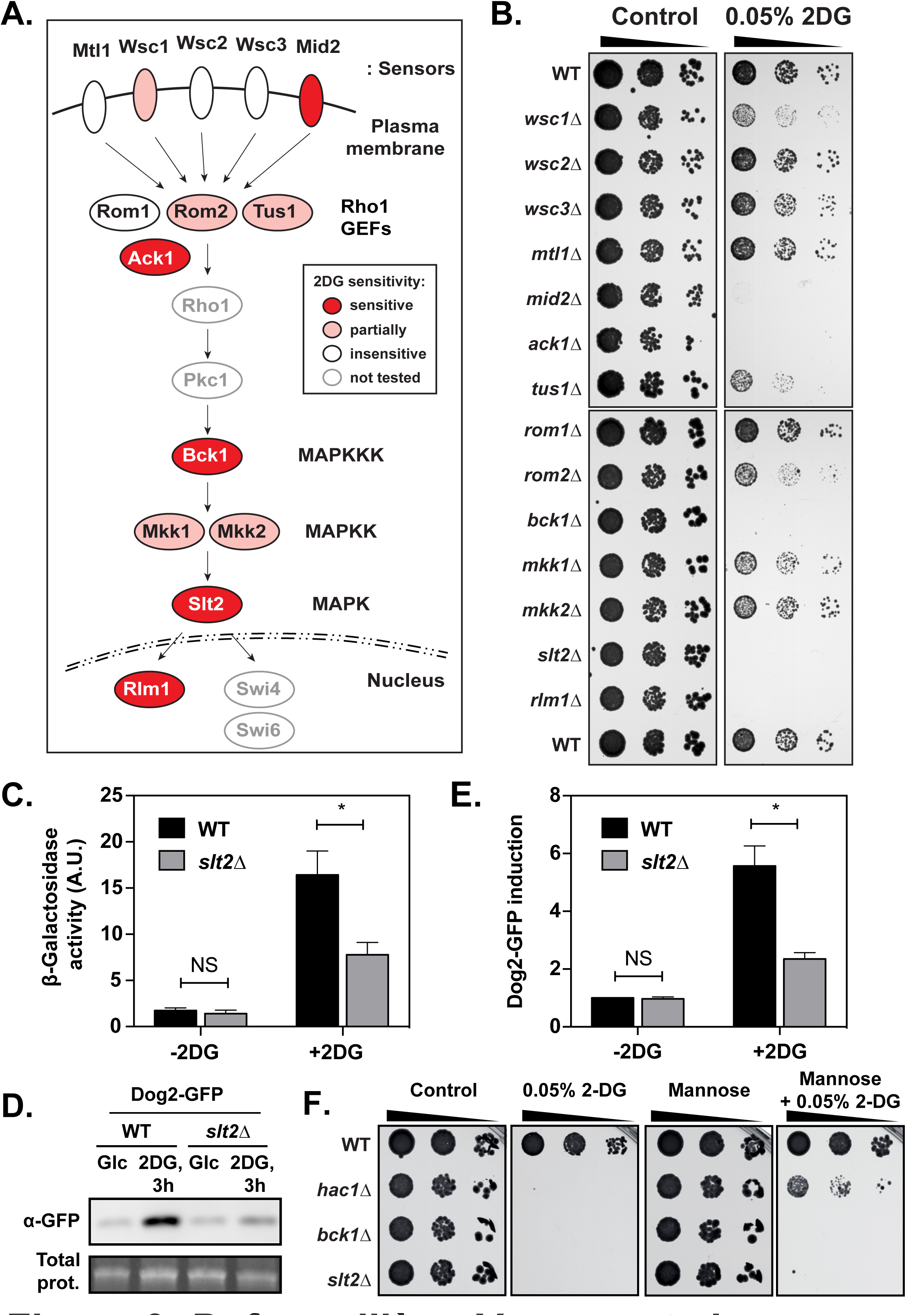
2DG activates the MAPK-based cell-wall integrity pathway, which is required for 2DG tolerance and additionally contributes to the regulation of Dog2 expression. (**A**) Schematic of the cell wall integrity pathway showing the various actors and their requirement for growth on 2DG (see color code in the inset) based on drop tests showed in panel (**B**). (**B**) Serial dilutions of cultures from the indicated deletion strains were spotted onto SD plates containing no DG or 0.05% 2DG, and grown for 3 days at 30°C. (**C**) Beta-galactosidase assays on WT and *slt2∆* cells expressing *LacZ* under the control the *DOG2* promoter, before and after 3h 2DG treatments (± SEM, *n*=3). (**D**) Western blot on total protein extracts of WT and *slt2∆* cells expressing an endogenously tagged Dog2-GFP fusion, before and after 3h treatment with 2DG, using an anti-GFP antibody. (**E**) Relative expression of Dog2-GFP in the same conditions as (**D**) after normalization by total proteins and using WT/untreated as a reference (± SEM, *n*=3). (**F**) Serial dilutions of cultures from the indicated strains were spotted onto SC plates (supplemented with 2% mannose when indicated) containing no DG or 0.05% 2DG, and were grown for 3 days at 30°C.

We thus questioned whether CWI activation by 2DG contributed to Dog1 and Dog2 induction. Indeed, the activity of both promoters was decreased in the *slt2∆* mutant (Figure 3C and S4C). This was confirmed for Dog2 by analyzing the expression levels of Dog2-GFP (Figure 3D-E). Thus, similarly to tunicamycin (*59*), 2DG induces cell-wall integrity signaling and participates in Dog1 and Dog2 induction, in addition to the UPR pathway. Interestingly, however, the addition of exogenous mannose did not lead to a better tolerance of CWI mutants to 2DG (Figure 3F). Thus, ER stress relief is not sufficient to suppress 2DG toxicity in these mutants, suggesting that 2DG has other effects in the cell beyond triggering ER stress.

### A third pathway negatively regulates the expression of Dog2, but not Dog1, by glucose availability and participates in the described resistance of glucose-repression mutants to 2DG

The toxicity of 2DG also lies in the fact that it impairs glycolysis by inhibiting hexokinase and phosphoglucose isomerase activity (*6–8*). This leads to an energetic stress and accordingly, 2DG treatment activates AMPK in mammals (*62*).

Several lines of evidence indicate that the activity of the yeast AMPK orthologue, Snf1, is important for 2DG tolerance. Whereas the *snf1∆* mutant is hypersensitive to 2DG, the *reg1∆* mutant, in which Snf1 is hyperactive, displays an increased resistance towards 2DG which depends on Snf1 activity (*31*). We then questioned whether some of the resistance/sensitivity phenotypes associated to this pathway (Figure 4A) could be due to an altered level of Dog1 or Dog2 expression, particularly in the light of a previous study showing that Snf1 positively regulates Dog2 expression (*48*). To confirm the effect of Snf1 on Dog1 and Dog2 expression, glucose-grown cells were transferred to a medium containing lactate as a sole carbon source, which is known to cause Snf1 activation and consequently, the de-repression of glucose-repressed genes (Figure 4A). The expression of Dog2 was increased in this condition, but not that of Dog1 and this occurred in a Snf1-dependent manner (Figure 4B). In line with these data, the *pDOG2*-LacZ reporter was induced after transfer to lactate medium, but this response was decreased in the *snf1∆* mutant, confirming that *DOG2* is subject to a glucose-repression mechanism and is thus constantly repressed in a *snf1∆* mutant (Figure 4C), which may explain why the *snf1∆* mutant is sensitive to 2DG. A previous report indicated that Dog1/2 overexpression in a *snf1∆* mutant did not rescue resistance to 2DG, but we thought that this might be due to the fact that these experiments involved a high-copy plasmid in which Dog2 expression was still under the control of its endogenous, i.e. Snf1-regulated, promoter (*47*). Indeed, when overexpressed using a strong and constitutive promoter (*pGPD*), Dog2 could rescue Snf1 growth on 2DG (Figure 4D).

**Figure 4.**
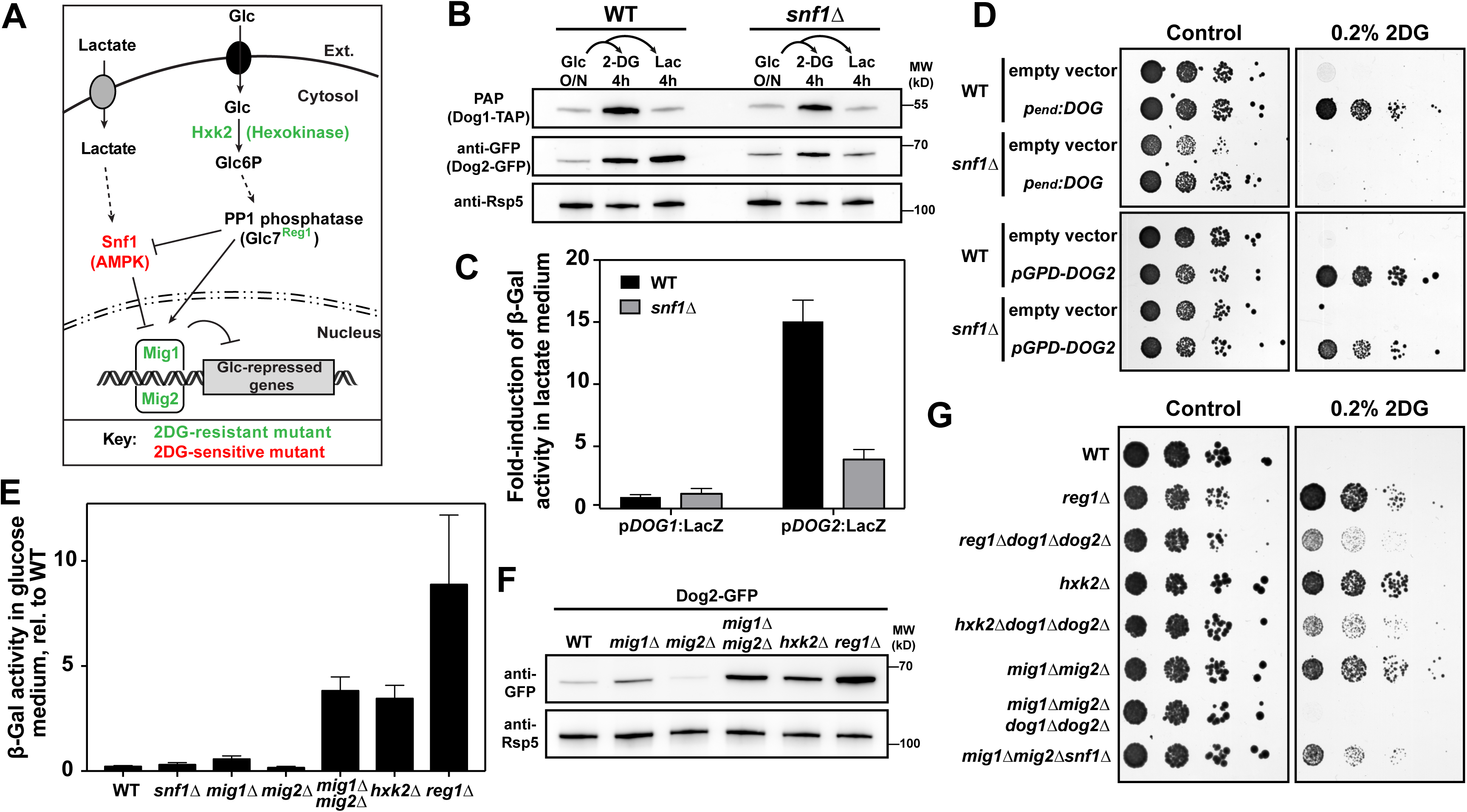
*DOG2*, but not *DOG1*, is controlled by glucose availability through transcriptional repression by Mig1-Mig2 and the kinase Snf1. (**A**) Schematic of the glucose repression pathway showing how Snf1/AMPK, PP1 (Glc7/Reg1) and their downstream transcriptional repressors Mig1/Mig2 regulate glucose-repressed genes in response to glucose availability or absence (eg. lactate). Green: 2DG-resistant mutants, red: 2DG sensitive mutant (see Introduction and panel **G**). (**B**) WT and *snf1∆* strains, both expressing endogenously tagged Dog1-TAP and Dog2-GFP fusions, were grown overnight in SC medium and then either treated with 0.2% 2DG or switched to an SC-lactate medium for 4 hours. Dog1-TAP was detected with the peroxidase-anti-peroxidase (PAP) complex, Dog2-GFP with anti-GFP antibodies. The anti-Rsp5 antibody was used to control loading. (**C**) Beta-galactosidase activity on WT and *snf1∆* cells expressing *LacZ* under the control the *DOG1* or the *DOG2* promoter, before and after 3h growth in lactate. The fold-induction after transfer to lactate is indicated for each promoter in each strain (± SEM, *n*=3). (**D**) WT and *snf1∆* strains, transformed with either a genomic clone containing both *DOG1* and *DOG2* under the control of their own promoter (*pend:DOG*, top panels), or with a vector containing *DOG2* under the control of the strong *GPD* promoter (*pGPD:DOG2*, bottom panels) were grown, serial-diluted and spotted onto SC plates (SC-Leu, top plates, and SC-Ura, bottom plates) with or without 0.2% DG, and grown for 3 days at 30°C. (**E**) Beta-galactosidase assays on WT and the indicated deletion mutants expressing *LacZ* under the control the *DOG2* promoter after overnight growth in SC medium (exponential phase) (± SEM, *n*=3). (**F**) The indicated strains, all expressing an endogenously tagged Dog2-GFP fusion, were grown overnight in SC medium (exponential phase). Total protein extracts were prepared and blotted with anti-GFP antibodies and the anti-Rsp5 antibody was used as a loading control. (**G**) Serial dilutions of cultures of the indicated mutants were spotted on YPD plates containing 0, 0.2% or 0.5% of 2DG. Plates were scanned after 3 days of growth at 30°C.

In contrast, Dog2 expression was increased in glucose-repression mutants which display an increased Snf1 activity, such as mutants lacking the hexokinase Hxk2 or the PP1 regulatory phosphatase Reg1 (*44, 63*), both at the promoter level and the protein level (Figure 4E-F). In addition, lack of the Snf1-regulated transcriptional repressors Mig1/Mig2 (see Figure 4A) also led to an increase in Dog2 expression (Figure 4E-F). A regulatory sequence in the promoter, present between −250 and - 350 bp relative to the ATG, combined with a proposed Mig1-binding site located at ca. −200 bp (*40, 48*) was critical for this glucose-mediated repression (Figure S5). Interestingly, mutants showing an increased Dog2 expression such as *reg1∆* or *hxk2∆* are known to be more tolerant to 2DG than the WT (*30, 31*), and we observed that this is also the case for the double mutant *mig1∆ mig2∆* (Figure 4G). We deleted *DOG1* and *DOG2* in these mutants to evaluate their contribution to 2DG resistance. The absence of Dog1 and Dog2 sensitized all strains to 2DG (Figure 4G); in particular, the *mig1∆ mig2∆ dog1∆ dog2∆* regained a sensitivity that is comparable to that of the WT, demonstrating that the resistance displayed by the *mig1∆ mig2∆* mutant is due to an increased *DOG* expression. In contrast, the *reg1∆ dog1∆ dog2∆* and the *hxk2∆ dog1∆ dog2∆* strains still remained more resistant than the WT, suggesting additional mechanisms of resistance. Finally, we observed that the *snf1∆ mig1∆ mig2∆* was resistant to 2DG despite the absence of Snf1, in line with the idea that the *snf1∆* is 2DG-sensitive because of the constitutive repression of Mig1/Mig2 target genes, such as Dog2 and possibly other genes (Figure 4F). Overall, we conclude that Dog2 presents an additional regulation by glucose availability through Snf1/AMPK activity, which participates in the well-described resistance of glucose-repression mutants to 2DG.

### Increased expression of *DOG2* is frequently observed in spontaneous 2DG-resistant clones

Previous screens led to the identification of 2DG-resistant mutants (*26, 29, 35*). The initial purpose of these screens was to identify mutants that were insensitive to the repressive effect of 2DG, or that were impaired for glucose phosphorylation (since only 2DG6P is toxic to cells), and consequently, were performed on media containing other carbon sources than glucose.

The mechanisms involved in 2DG resistance on glucose medium were tackled much later by screening of the deletion library (*30*), which led to the identification of resistant mutants, some of which were subsequently confirmed (*31*). In the course of our experiments, we often found that 2DG-resistant clones could spontaneously arise from wild-type or even 2DG-susceptible strains (see for example Figure 1E). It was previously noted that yeast acquires 2DG resistance at a high frequency, which can be accompanied by an increase in 2DG-6-phosphatase activity (*28, 38*). To study whether Dog2, which is functionally the most important paralogue for 2DG resistance, is upregulated during the emergence of spontaneous 2DG-resistant mutants, we spread ca. 2×10^6^ cells on a glucose-based medium containing 0.2% 2DG, and selected clones that had appeared after 6 days of growth. Altogether, 24 clones were obtained, whose resistance to 2DG was confirmed (see Figure 5B). Using the p*DOG2*-LacZ reporter, we found that 13 clones displayed a significantly increased *DOG2* promoter activity as compared to WT strains (Figure 5A). This suggested that Dog2 overexpression is a frequent feature of 2DG-resistant clones. Among those, we expected to isolate *reg1* and *hxk2* mutants because both are resistant to 2DG in glucose medium (See Figure 5G; *30, 31*). To identify them, we first tested whether some of the isolated resistant clones displayed phenotypes typical of *reg1* mutants, such as sensitivity to tunicamycin (*64, 65*) or selenite (which enters the cell through the glucose-repressed transporter Jen1: *66, 67*), which allowed to discriminate potential *reg1* mutants from *hxk2* mutants (Figure 5B). Indeed, three of the isolated clones were potential *reg1* mutants based on these phenotypes (Figure 5B). This was further supported by the observation that these 3 mutants displayed a markedly increased activation of Snf1, as measured by using an antibody directed against phosphorylated AMPK (Figure S6A). Sequencing of the *REG1* coding sequence in these clones revealed point mutations (substitutions) causing the appearance of stop codons which likely explain Reg1 loss of function (Figure 5C). To additionally identify whether the set of spontaneous resistant clones contained *hxk2* mutants, we performed a complementation test and therefore crossed these strains with either a WT strain or an *hxk2∆* strain and tested the ability of the resulting diploids to grow on 2DG. All diploids generated through the cross with the WT lost their ability to grow on 2DG, except two (clones #23 and #24) (Figure 5D), indicating that 2DG resistance was generally caused by (a) recessive mutation(s). In contrast, 12 diploids obtained by the cross with the *hxk2∆* strain maintained their ability to grow on 2DG (Figure 5D), suggesting that 2DG resistance of the initial strains was caused by a deficiency in *HXK2* function. Most of these clones also displayed an increased p*DOG2*-LacZ expression, in agreement with our previous findings (Figure 4E-F). We PCR-amplified and sequenced the *HXK2* ORF in the WT and these mutants (Figure 5E). Out of the 12 mutants identified that are in the same complementation group as *HXK2*, two did not display any mutation in the *HXK2* ORF (clones #1 and #8) and may represent regulatory mutants in *cis*, possibly in regions that were not sequenced. In favor of this hypothesis, we found that their 2DG sensitivity can be restored by re-expression of *HXK2* using a multicopy, genomic clone (Figure S6B). The other 10 mutants carried at least one mutation in the *HXK2* coding sequence (Figure 5E). Four mutations created a stop codon (clones #12, #15, #16 and #20) resulting in a predicted protein that was truncated almost fully (#20: mutation Leu^4^>TAA) or by more than half of its length, which is expected to lead to loss of function since conserved regions present in the C-terminal domain are essential for ATP binding and hexokinase activity (*68, 69*). Five mutants (#2, #3, #11, #13 and #14) carried a single point mutation in residues conserved across all hexokinase sequences tested (see alignment, Figure S6, C-D). Four of these affected residues in close proximity to glucose binding residues (#2: T^212^>P; #3: K^176^>T; #11 and #14: Q^299^>H) (*70*). The fifth mutation (#13: G^418^>C) was located next to S^419^, proposed to interact with the adenosine moiety of ATP (*68, 70*). Finally, one mutant (#21) bared two mutations, T^75^>I and S^345^>P, the latter of which introduces a Pro residue nearby a putative ADP-binding pocket (*70, 71*). However, these residues are much less conserved (Figure S6D) so it could be that both mutations combined are required to abolish Hxk2 function, although this is only speculative.

**Figure 5.**
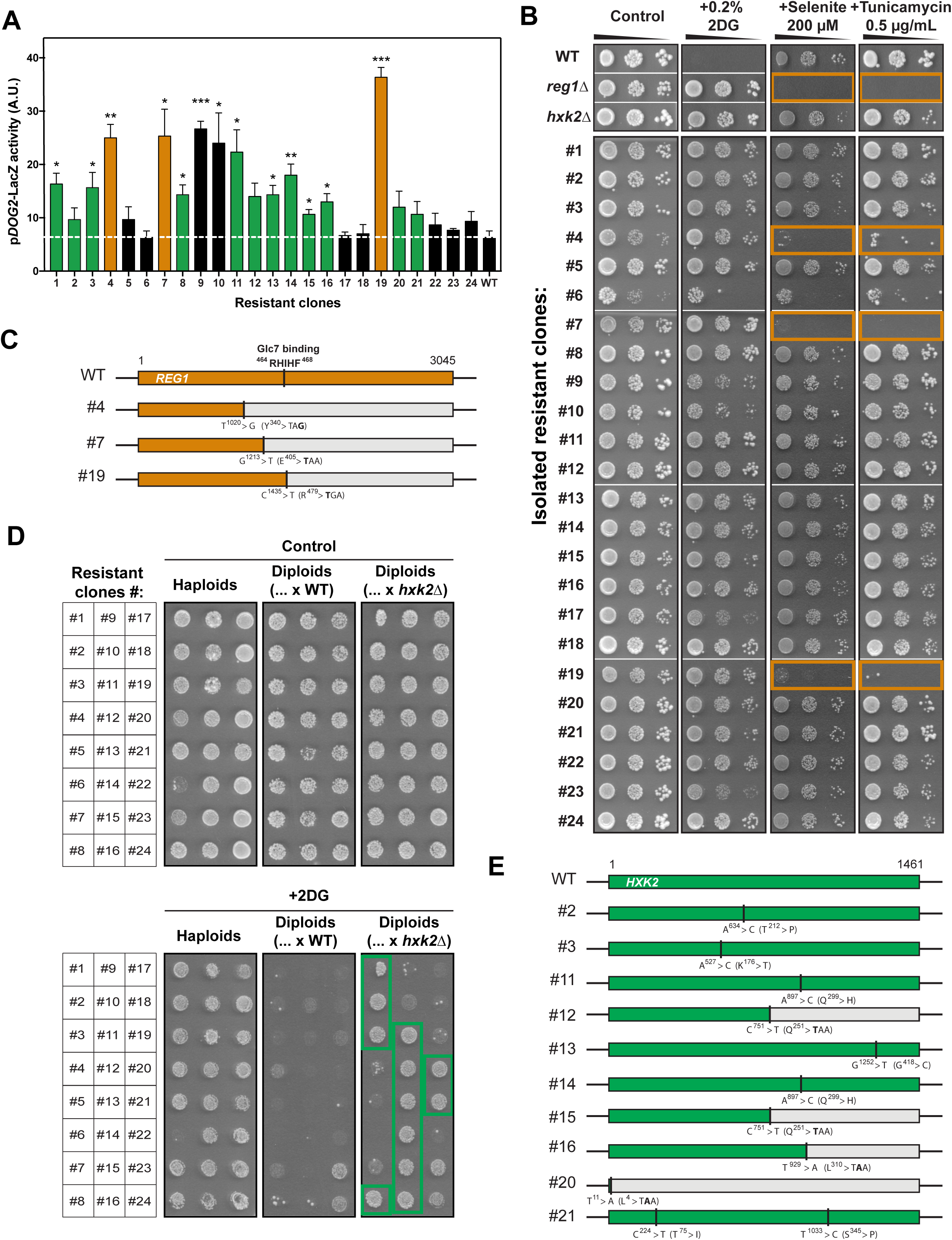
Most clones that become spontaneously resistant to 2DG show an increased Dog2 expression. (**A**) 24 clones showing a spontaneous resistance to 0.2% 2DG were isolated. The beta-galactosidase activity of these mutants, due to the expression of the *LacZ* reporter driven by the *DOG2* promoter, was measured after overnight growth in SC medium (exponential phase) (± SEM, *n*=3). For colors, see panels (**C**) and (**E**). (**B**) Serial dilutions of cultures from the indicated deletion strains and resistant clones were spotted onto SC plates containing no DG or 0.2% 2DG, as well as 200 µM selenite or 0.5 µg/mL tunicamycin and grown for 3 days at 30°C. Note that clone #6 grows very slowly even in absence of 2DG. Orange squares: identified *reg1* mutants. (**C**) Schematic of the identified mutations within the *REG1* ORF in the indicated mutants as obtained by sequencing. (**D**) The 24 resistant clones were crossed with a WT or an *hxk2∆* strain of the opposite mating type, and cultures of the haploid resistant clones (see matrix*, left*) or the resulting diploids were spotted onto SC medium with or without 0.2% DG. Green squares: identified *hxk2* mutants. (**E**) Schematic of the identified mutations within the *HXK2* ORF in the indicated mutants as obtained by sequencing.

Among the 2DG-resistant clones which expressed *pDOG2*-LacZ at a high level, 2 clones (#9 and #10) remained unaccounted for by mutations in either *REG1* or *HXK2*, suggesting that their resistance involved a novel mechanism. Whole genome resequencing of the genomic DNA isolated from these clones and comparison with that of the parent strain revealed several SNPs (**Table S2**) including a nonsense mutation in *CYC8* (C^958^>T) that was identified in both strains, causing a premature stop codon at position 320 (Figure 6A). The *CYC8* gene is also known as *SSN6* (*Suppressor of snf1*) and mutations in this gene lead to constitutive expression of the glucose-repressed gene encoding invertase (*SUC2*), even in a *snf1* mutant (*72, 73*). Indeed, *CYC8* encodes a transcriptional co-repressor which controls the expression of glucose-regulated genes (*74*). Because the repression of *DOG2* expression by glucose is controlled by Snf1 and Mig1/Mig2 (see Figure 4), we hypothesized that *CYC8* could also take part in *DOG2* regulation, so that a mutation in *CYC8* could lead to increased 2DG resistance through *DOG2* overexpression. We observed that indeed, mutants #9 and #10 expressed invertase even in when grown in glucose medium (repressive conditions), in agreement with a mutation in *CYC8* (Figure 6B). We also observed that these mutants displayed an increased adhesion phenotype, i.e. persistence of colony structures upon washing the plate (*75*) that is often associated to mutants that are prone to flocculate, such as *cyc8* mutants (*76, 77*) (Figure 6C). When we introduced a low copy (centromeric) plasmid containing *CYC8* under the control of its endogenous promoter, we observed that it suppressed the 2DG resistance of mutants #9 and #10 as well as their adhesion to the agar plate (Figure 6C). This was not the case when using a truncated version of *CYC8* identified in this screen (Figure 6C), although mutants #9 and and #10 did not display a slow growth contrary to the *cyc8∆* mutant, suggesting that it is at least partially active for other functions. Finally, we tested whether the strong activity of the *DOG2* promoter observed in mutants #9 and #10 (see Figure 5A) was also due to the lack of a functional *CYC8*. Indeed, the re-introduction of low-copy vector containing *CYC8* led to a decreased reporter expression in these mutants (Figure 6D). This was confirmed by examining Dog2-GFP expression at the protein level (Figure 6E-F). Therefore, *DOG2* expression is regulated by Cyc8 and spontaneous *cyc8* mutants display an increase in Dog2 expression and increased resistance to 2DG.

**Figure 6.**
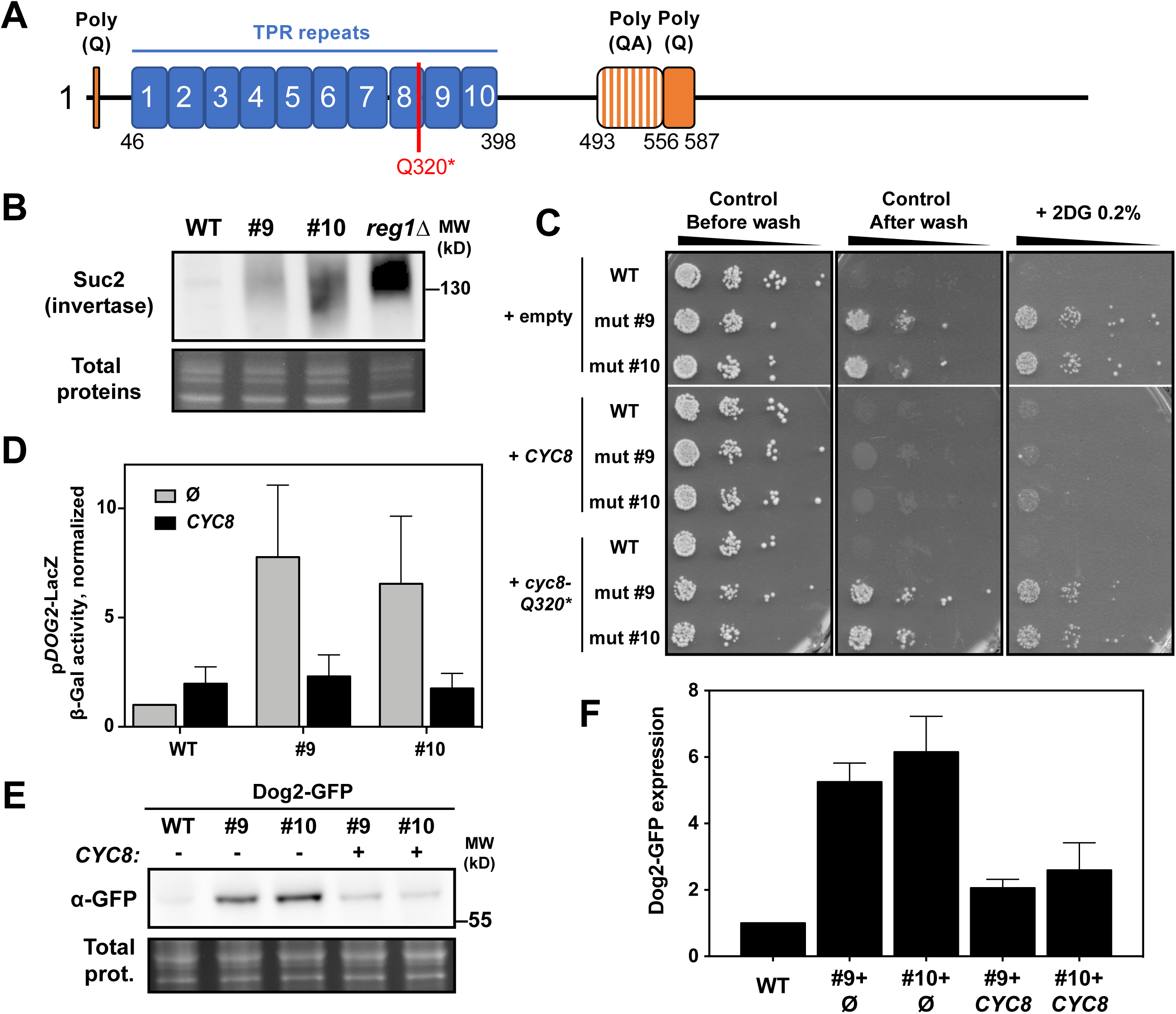
The 2DG-resistant phenotype of mutants #9 and #10 is caused by a mutation in *CYC8*. (**A**) Schematic of the domain organization of the Cyc8 protein, showing the Poly(Q) and Poly(QA) repeats as well as the N-terminal TPR repeats. Red: mutation identified by whole genome resequencing of the spontaneous 2DG-resistant mutants #9 and #10. (**B**) A WT strain, the mutant strains #9 and #10 and the *reg1∆* mutant (positive control) were grown in SC medium (exponential phase). Total protein extracts were prepared and blotted using anti-invertase antibodies (invertase is heavily glycosylated and migrates as a smear (*117*)). (**C**) WT and mutants #9 and #10 were transformed with low-copy (centromeric) plasmid either empty or containing WT *CYC8* or mutant *cyc8 (Q320*)*, and were spotted on SC-Leu or SC-Leu + 0.2% 2DG medium, and grown for 3 days at 30°C. Middle panel: the control plate was scanned and then washed for 1 min under a constant flow of water, and then scanned again (see Methods). (**D**) Beta-galactosidase activity on WT and mutants #9 and #10 expressing *LacZ* under the control the *DOG2* promoter and transformed with an empty vector or a low-copy vector containing WT *CYC8*, after growth in SC medium (normalized to the value of the WT, ± SEM, *n*=3). (**E**) Western blot on total protein extracts of WT and mutants #9 and #10 cells expressing an endogenously tagged Dog2-GFP fusion and transformed with either an empty plasmid or a low-copy (centromeric) plasmid containing WT *CYC8* after growth in SC medium, using an anti-GFP antibody. (**F**) Relative expression of Dog2-GFP in the same conditions as (**G**) after normalization by total proteins and using the WT control as a reference (± SEM, *n*=3).

Altogether, this initial characterization revealed that *DOG2* overexpression is a common phenomenon within spontaneous 2DG-resistant clones, both in known 2DG-resistant mutants (*reg1* and *hxk2*) and in the novel 2DG-resistant *cyc8* mutants that we isolated.

### HDHD1, a human member of the HAD-like phosphatase family, is a new 2DG-6-P phosphatase involved in 2DG resistance

The fact that Dog2 expression is increased in various spontaneous 2DG-resistant mutants reminded us of early studies in HeLa cells showing an increased 2DG6P phosphatase activity in isolated 2DG resistant clones (*20*). Dog1 and Dog2 belong to the family of HAD (Haloacid Dehalogenase)-like phosphatases conserved from bacteria to human (Figure 7A). The bacterial homolog of Dog1/Dog2, named YniC, can also dephosphorylate 2DG6P *in vitro* (*78*) and we found that the expression of YniC in the double *dog1∆ dog2∆* yeast mutant also restored 2DG resistance (Figure 7B). We used this phenotype to identify potential human orthologues (Figure 7B). Running a PSI-BLAST (*79*) on the human proteome using the Dog2 protein sequence retrieved HDHD1-isoform a (NP_036212.3) as the best hit (39% homology). HDHD1 (for Haloacid Dehalogenase-like Hydrolase Domain containing 1; also named PUDP for pseudouridine-5’-phosphatase) is a HAD-like phosphatase with a demonstrated *in vitro* activity towards phosphorylated metabolites such as pseudouridine-5’-phosphate (*80*). When expressed in in the double *dog1∆ dog2∆* mutant, HDHD1 was partially able to rescue growth on 2DG containing medium (Figure 7B). The expression of HDHD4 (also named NANP, for N-acylneuraminate-9-phosphatase; *81*), which belongs to the same subfamily as HDHD1 within the HAD-phosphatase family (37% homology), did not significantly restore yeast growth on 2DG, neither did that of PSPH (phosphoserine phosphatase; *82*), another close family member (Figure S7A,B). Moreover, among the four predicted isoforms of HDHD1 (isoforms 1-4), only HDHD1-1 rescued the growth of the *dog1∆ dog2∆* on 2DG-containing medium (Figure S7C) despite the fact that all isoforms were expressed in yeast, as confirmed by western blotting using anti-HDHD1 antibodies (Figure S7D).

**Figure 7.**
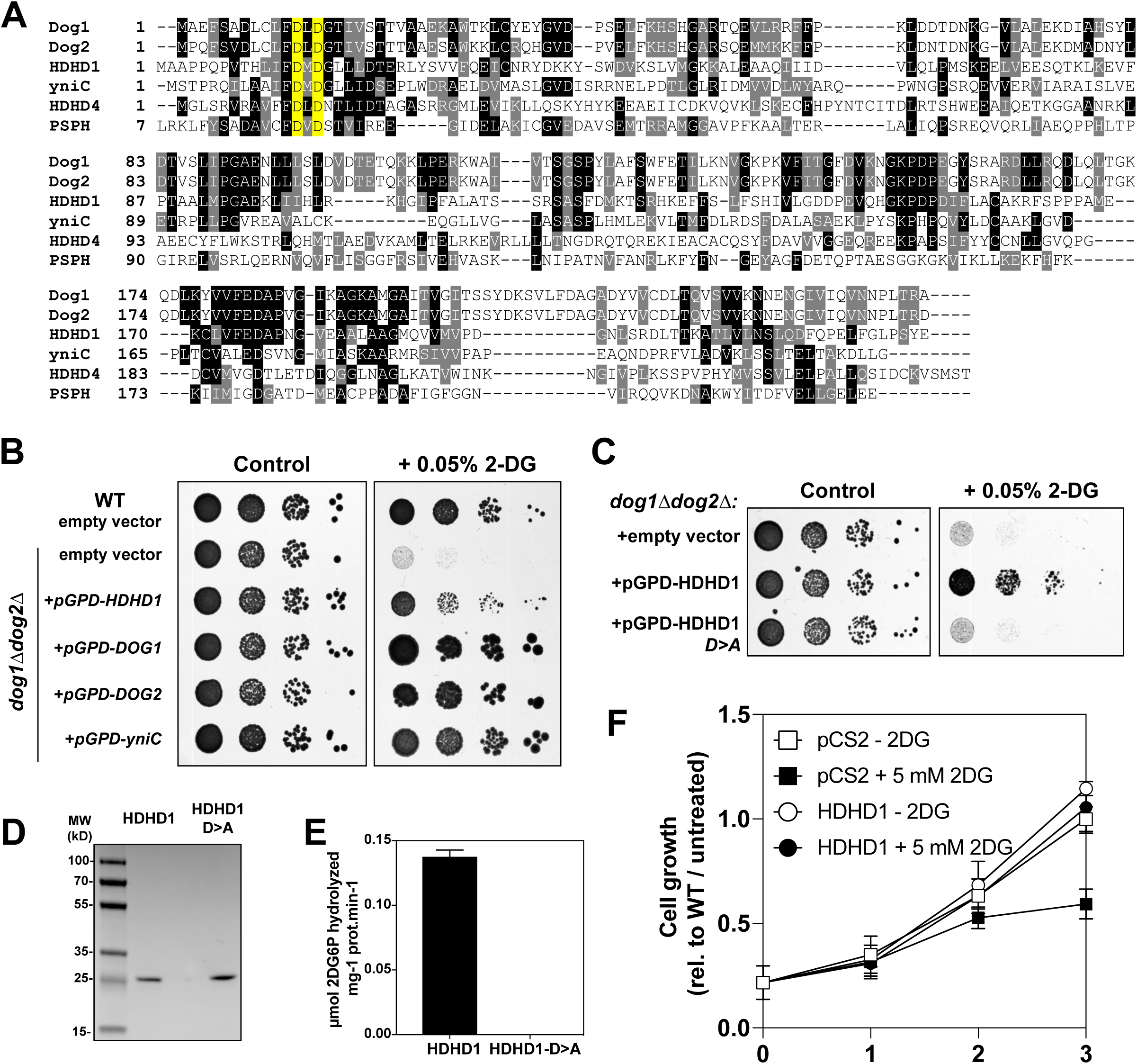
The human phosphatase HDHD1 has a 2DG6P phosphatase activity and its overexpression leads to 2DG resistance in HeLa cells. (**A**) Multiple protein sequence alignment of yeast Dog1, Dog2, *Escherichia* coli yniC and the human proteins HDHD1, HDHD4 and PSPH aligned with ClustalX 2.0. The highly conserved catalytic aspartates are displayed in yellow. The six first amino-acids of PSPH were truncated to optimize the N-terminal alignment of its catalytic aspartates with the other phosphatases. (**B**) Serial dilutions of WT and *dog1∆dog2∆* strains transformed with the indicated plasmids were spotted on SC-Ura medium with or without 0.05% 2DG and were scanned after 3 days of growth at 30°C. (**C**) Serial dilutions of *dog1∆dog2∆* strains transformed with an empty vector or vectors allowing the expression of a HDHD1 or its predicted catalytic mutant, HDHD1-DD>AA (mutations of the N-terminal catalytic aspartates to alanines) were spotted on SC-Ura medium with or without 0.05% 2DG and were scanned after 3 days of growth at 30°C. (**D**) Recombinant, His-tagged HDHD1 and HDHD1-DD>AA were expressed in bacteria and purified for in vitro enzymatic tests. 0.7µg were loaded on a gel to show homogeneity of the protein purification. (**E**) *In vitro* 2DG6P phosphatase activity of HDHD1 and HDHD1>DDAA as measured by assaying glucose release from 2DG6P. (**F**) Growth of HeLa cells transfected with an empty vector (□) or with a construct allowing the overexpression of HDHD1 (○) over time in the absence (open symbols) or presence (filled symbol) of 5mM 2DG. The number of cells is normalized to that of the untransformed/untreated cells after 3 days (± SEM, *n*=3).

The mutation of conserved aspartate residues (D^12^ and D^14^, see alignment in Figure 7A), predicted to be essential for the catalytic activity of HAD phosphatases (*83*) abolished the ability of HDHD1 to restore growth of the *dog1∆ dog2∆* mutant on 2DG (Figure 7C and S7E), suggesting it may act as a 2DG-6P phosphatase. This was confirmed by purifying recombinant HDHD1 (Figure 7D) and testing for a potential 2DG-6-P phosphatase activity. Indeed, HDHD1 could dephosphorylate 2DG-6P *in vitro*, and this activity depended on the integrity of its putative catalytic residues (Figure 7E). We then tested whether HDHD1 overexpression in HeLa cells could lead to an increased resistance to 2-DG. Low concentrations of 2DG (5 mM) in the presence of glucose (25 mM) were sufficient to inhibit the growth of cells transfected with an empty vector, whereas those that overexpressed HDHD1 were virtually insensitive to 2DG treatment (Figure 7F). These results suggest that a dysregulated expression of HDHD1 could modulate 2DG resistance in human cells.

## Discussion

The first studies examining the effect of 2DG on glycolysis in yeast but also in normal and cancer tissues were published more than 60 years ago (*84–86*). Yet, its mode of action is not fully understood. Data suggests that 2DG can be considered as a general competitor of glucose (*87*) and as a glycolysis inhibitor because of its inhibition of glycolytic enzymes (*6–8*). However, this view has been challenged since exposure of cancer cells to 2DG interferes with N-linked glycosylation, likely because of the structural similarity of 2DG with mannose (*88*). This results in ER stress and UPR induction in mammalian cells (*14, 16*). Therefore, 2DG interferes with other cellular functions beyond glycolysis.

In this study, we used mass spectrometry as an unbiased approach to describe the cellular effects of 2DG on the total cellular proteome. This revealed that many glycolytic enzymes are upregulated, likely as a consequence of an impaired glycolysis, as well as many genes regulated by the MAPK-based cell wall integrity pathway, suggestive of its activation (discussed below).

A second and unexpected observation was to identify, within the list of 2DG-upregulated proteins, the Dog1 and/or Dog2 phosphatase previously involved in 2DG resistance (*33, 40, 41*). Dog2 upregulation upon 2DG treatment was intriguing because it wasn’t clear how a synthetic molecule such as 2DG could trigger the onset of a resistance mechanism. It should be noted that the cellular functions of Dog1/Dog2, beyond 2DG dephosphorylation, are unknown. They can dephosphorylate a range of phosphorylated sugars *in vitro* (*40*), in agreement with the fact that other members of the HAD family of phosphatases can accommodate various substrates (*78*). Therefore, it seemed plausible that Dog1/2 induction was a response to one or more of the cellular consequences of 2DG treatment, and not to 2DG itself. Our study reveals that actually, the expression of Dog1 and Dog2 is controlled by multiple signaling pathways, each of which are turned on as a response to 2DG (see Working model, **Figure 9**). This study was also taken as an opportunity to probe the signaling pathways that are activated upon 2DG exposure and precise the causes of these activations.

**Figure 8.**
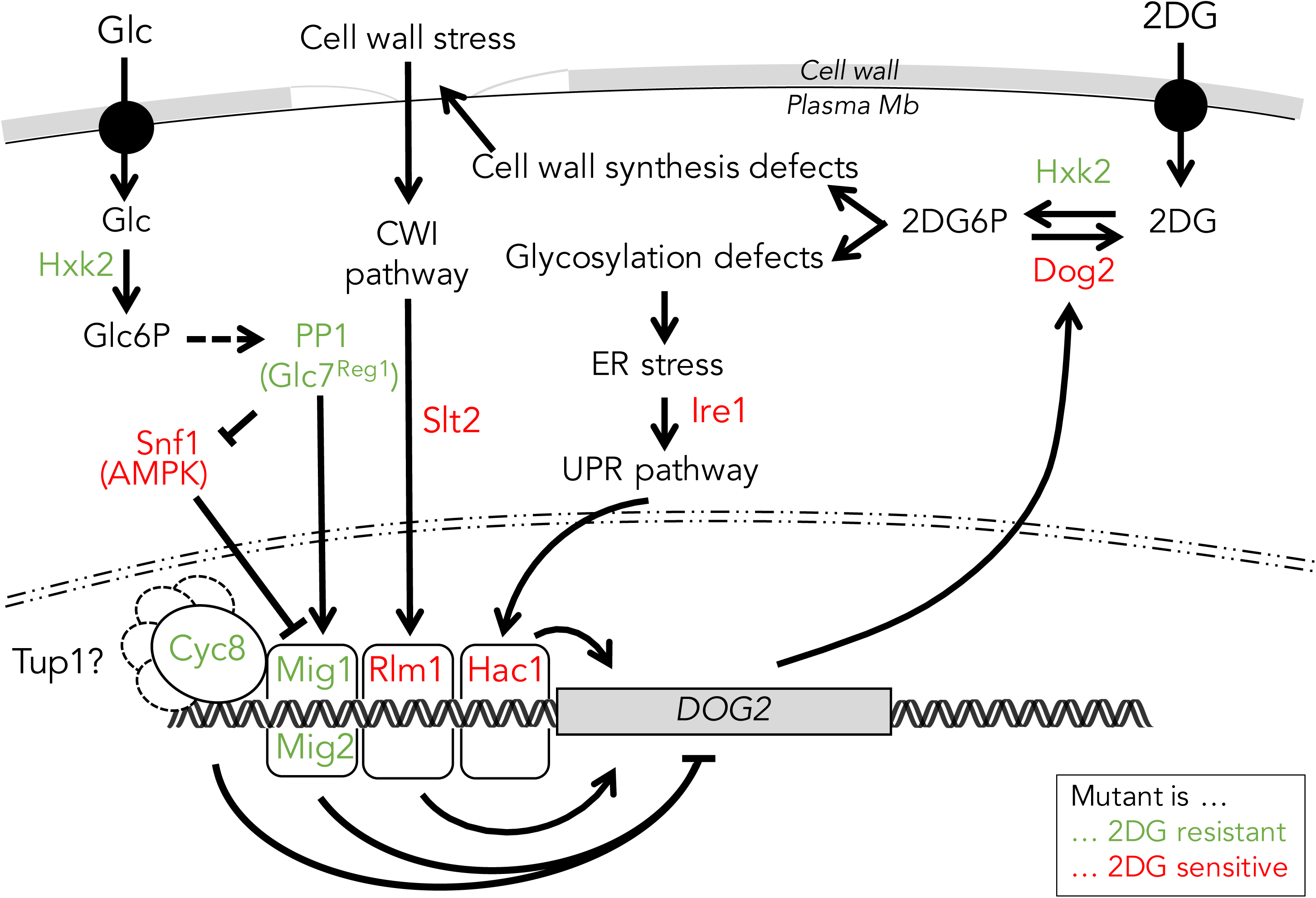
Working Model. Glucose phosphorylation triggers the onset of the glucose-repression pathway, by which PP1 inactivates Snf1/AMPK. This leads to the lack of phosphorylation of Mig1/Mig2, which remain in the nucleus to mediate the glucose-repression of genes such as *DOG2*. The deletion of *REG1*, *HXK2*, or *MIG1* and *MIG2*, or a mutation in *CYC8* leads to 2DG resistance, at least in part through an increased expression of *DOG2* which dephosphorylates 2DG-6-P. Instead, the deletion of *SNF1* causes an increased sensitivity to 2DG, which can be rescued by the deletion of *MIG1* and *MIG2* or by Dog2 overexpression. In parallel, 2DG6P causes (i) ER stress and triggers the UPR pathway, which stimulates *DOG2* expression through the transcription factor Hac1, and (ii) the CWI integrity pathway, likely through interference with polysaccharide and cell wall synthesis, which also induces *DOG2* through the transcription factor Rlm1.

First, we confirmed that 2DG triggers ER stress and the onset of the UPR pathway. This was not unexpected given the proposed mode of action of this drug and the available data in the literature on mammalian cells (*14, 16*), but had not been formally proven in yeast with current tools. The induction of an UPR reporter upon 2DG treatment and the hypersensitivity of UPR mutant strains such as *hac1∆* or *ire1∆* support this conclusion. The latter phenotype could be alleviated by restoring N-glycosylation through the addition of mannose in the culture medium. We also showed that Dog1 and Dog2 are induced by ER stress, in line with high-throughput studies suggesting that Dog1 and Dog2 are UPR-regulated genes (*89*).

Second, we showed that 2DG activates the MAP kinase-based cell wall integrity (CWI) pathway, which is in charge of the cellular response to cell wall alteration and other stresses. Several studies reported connections between ER stress signaling and the CWI pathway (*57–61*), and 2DG-elicited N-glycosylation and ER stress may have repercussions on the CWI pathway. For instance, CWI pathway mutants are sensitive to tunicamycin because they lack an ER surveillance-pathway which normally prevents the inheritance of stressed ER to the daughter cell during cell division (*59*). Many CWI mutants were indeed sensitive to low concentrations (0.05%) of 2DG but unlike UPR mutants, their growth was not restored by the addition of exogenous mannose, suggesting that N-glycosylation defect is not the sole reason why CWI mutants are hypersensitive to 2DG. However, 2DG interferes with the synthesis of structural polysaccharides that make up the yeast cell wall, again because 2DG acts as an antagonist of mannose and glucose incorporation into these polymers (*42, 90–92*). 2DG exposure leads to yeast cell lysis at sites of growth, which is where glucan synthesis occurs (*43*) and where the major glucan synthase, Fks1, is localized (*93*). 2DG-induced weakening of the cell wall could trigger the cell wall integrity pathway (**Figure 9**). This would explain why the main sensors responsible for sensing cell wall damage, Wsc1 and Mid2 (reviewed in *56*), are required for growth on 2DG. In addition, we report that Dog1 and Dog2 are less expressed in CWI mutants, which could further sensitize these strains to 2DG. The effect of 2DG on cell wall synthesis denotes an interference of 2DG with UDP-glucose metabolism that is likely to be conserved in metazoans (*94*), where it may affect metabolic pathways involving these precursors, such as glycogen synthesis (*95, 96*).

Our data also reveal that Dog2 expression is regulated at the transcriptional level by glucose availability through the glucose-repression pathway, which is controlled by the kinase Snf1/AMPK and the regulatory subunit of the PP1 phosphatase, Reg1. This explains why *reg1* mutants were shown to display an increased 2DG-6-phosphate phosphatase activity (*28, 38, 39*), since the glucose-mediated repression of Dog2 is defective in these mutants. The fact that Dog2 is regulated by this pathway may also contribute to the 2DG resistance of *hxk2* and *mig1* mutants, which participate in the glucose-repression pathways and were identified in previous screens (*29, 35*). Indeed, the 2DG resistance displayed by the *reg1∆* and *hxk2∆* strains was partially dependent on an increased expression of Dog2. In the *hxk2∆* mutant, the lack of hexokinase 2 is compensated by the expression of other glucose-phosphorylating enzymes (such as Hxk1 and glucokinase, Glk1) (*97*) that may be less prone to phosphorylate 2DG (*36*), and thus could lead to a lower accumulation of 2DG6P in the cell. Hxk2 also has additional, non-metabolic roles beyond sugar phosphorylation (reviewed in Ref. *98*), whose loss may additionally contribute to this phenotype. Yet, many of the *hxk2* point mutants isolated during the characterization of spontaneous 2DG-resistant clones were affected at or nearby glucose-binding residues, suggesting a primary metabolic role for Hxk2 in 2DG resistance. Concerning the *reg1∆* strain, additional mechanisms of resistance beyond an increased expression of Dog2 also remain to be investigated. What appeared from previous work is that the resistance of the *reg1∆* mutant is dependent on Snf1 hyperactivity, since the additional deletion of *SNF1* in this background restores 2DG sensitivity to the *reg1∆* strain (*31*). Thus, additional yet Snf1-dependent mechanisms cooperate with the increased expression of *DOG2* to allow *reg1∆* cells to resist to 2DG.

After 2DG import and phosphorylation, the amount of 2DG6P in cells can exceed that of G6P by up to 80-fold (*99*). Overexpression of Dog2 was previously shown to revert the repressive effect of 2DG (*33*), suggesting that it can clear the 2DG6P pool up to a point where 2DG6P is no longer detected by the yet unknown cellular glucose-sensing mechanism. Thus, Dog2 overexpression appears as a good strategy for 2DG resistance, and indeed, it was overexpressed in the majority of the spontaneous resistant mutants we isolated. This included mutants in the *REG1* and *HXK2* genes, but additionally, we identified two mutants carrying a point mutation in the gene encoding the transcriptional repressor Cyc8, leading to a truncated protein at codon 320, within its 8^th^ predicted tetratricopeptide (TPR) repeat. The TPR repeats are involved in the interaction of Cyc8 with its co-repressor Tup1, and to specify its recruitment to specific promoters, possibly through the recruitment of pathway-specific DNA-binding proteins (*100*). Point mutations in TPR units 9 and 10 affect Cyc8 function in a similar manner as the mutation we isolated (i.e. impact on glucose repression but not global growth), suggesting that alterations at this region only affect a subset of Cyc8 functions (*101*). This confirms the differential requirement of TPR repeats for the various functions of Cyc8, and in particular the involvement of the TPR repeats 8-10 in glucose repression (*102*). Altogether, these data describe *DOG2* overexpression as a successful strategy to overcome 2DG toxicity.

Similarly, the selection of HeLa-derived 2-DG-resistant cell lines showed an increase in 2DG6P phosphatase activity, but the responsible enzyme was not identified (*20*). Based on sequence similarity, we identified HDHD1 as an enzyme displaying *in vitro* 2DG6P phosphatase activity and whose overexpression in both yeast and HeLa cells allows resistance to 2DG. Actually, we found that HDHD1 has a very low affinity toward 2DG (in the 10^−2^ M range, unpublished data) as compared to other substrates such as 5ʹ-pseudouridine monophosphate or even 3ʹ-AMP (*80*). Despite this very low affinity for 2DG, HDHD1 overexpression in HeLa cells conferred resistance to 2DG. Whether HDHD1 is responsible for the 2DG resistance previously reported in human cells (*20*) remains to be investigated. Altogether, our work shows that 2DG-induced activation of multiple signaling pathways can rewire the expression of endogenous proteins which can target 2DG6P to promote 2DG tolerance, and whose deregulated expression can lead to 2DG resistance. This mechanism likely superimposes on other resistance mechanisms that should be scrutinized in the future.

## Material and Methods

### Yeast strain construction and growth conditions

All yeast strains used in this study derive from the *Saccharomyces cerevisiae* BY4741 or BY4742 background and are listed in **Supplemental Table S3**. Apart from the mutant strains obtained from the yeast deletion collection (Euroscarf) and the fluorescent GFP tagged strains originating from the yeast GFP clone collection (*103*), all yeast strains were constructed by transformation with the standard lithium acetate-polyethylene glycol protocol using homologous recombination and verified by PCR on genomic DNA prepared with a lithium acetate (200mM) / SDS (0.1%) method (*104*).

Yeast cells were grown in YPD medium (Yeast extract-Peptone-Dextrose 2%) or in synthetic complete medium (SC, containing 1.7g/L yeast nitrogen base (MP Biomedicals), 5g/L ammonium sulfate (Sigma-Aldrich), the appropriate drop-out amino acid preparations (MP Biomedicals) and 2% (w/vol) glucose unless otherwise indicated). Alternatively, SC medium could contain 0.5% lactate as a carbon source (from a 5% stock adjusted to pH=5; Sigma Aldrich). Pre-cultures of 4mL were incubated at 30°C for 8 hours and diluted in fresh medium on the evening to 20mL cultures grown overnight with inoculation optical densities (OD_600_) of 0.0003 for YPD and 0.001 for SC medium, giving a culture at mid-log phase the next morning.

For glucose depletion experiments, cultures were centrifuged and resuspended in an equal volume of SC/lactate medium and incubated at 30°C during the indicated times. For 2-deoxyglucose, NaCl and tunicamycin treatments, the compounds were added to mid-log phase yeast cultures grown overnight to respective final concentrations of 0.2% (w/v), 400mM and 1µg/mL and incubated for the indicated times. 2-deoxyglucose and tunicamycin were purchased from Sigma-Aldrich. The mannose-supplemented medium (Fig. 2B) consisted of an SC medium that contained all the element indicated above plus 2% (w/vol) mannose.

### Plasmid construction

All the plasmids presented in this study are listed in **Supplemental Table S4** and were directly constructed in yeast using plasmid homologous recombination (*105*). DNA inserts were amplified by PCR using 70-mer primers containing 50nt homology overhangs and Thermo Fisher Phusion High-fidelity DNA Polymerase and receiver plasmids were digested with restriction enzymes targeting the insertion region. Competent yeast cells rinsed with lithium acetate were incubated for 30 minutes at 30°C with 20µL of the PCR product and 1µL of the plasmid digestion product, followed by a heat shock at 42°C for 20 minutes and a recovery phase in rich medium (YPD) for 90 minutes at 30°C and plated on synthetic medium without uracil. The pDOG1/2-LacZ vectors were generated using the pJEN1-LacZ vector obtained from Bernard Guiard (*106*) and cloning the pDOG1 or pDOG2 promoters (1kb) at *Bgl*II and *Eco*RI sites.

The *DOG1*, *DOG2*, *HDHD1* (all four isoforms), *HDHD4*, *PSPH* and *yniC* overexpression vectors were obtained by digesting a pRS426 vector (2µ, *URA3*) containing the p*GPD* promoter with *Eco*RI and *Bam*HI enzymes. In parallel, inserts were PCR-amplified using the following DNA templates: *DOG1* and *DOG2* from yeast genomic DNA preparations (primers: oSL1166/1167 and oSL1141/1142), the *yniC* ORF from *Escherichia coli* DH5α cells (oSL1172/1173) and the human gene ORFs (Uniprot identifiers: HDHD1=Q08623-1/2/3/4, HDHD4=Q8TBE9 and PSPH=P78330) from DNA sequences generated by gene synthesis after codon optimization for yeast expression (Eurofins Genomics) (*HDHD1*: oSL1170/1171 for all isoforms except isoform 3: oSL1170/1216; *HDHD4*: oSL1214/1215; *PSPH*: oSL1212/1213). These PCR products were designed to include a 50-bp overlap with the digested plasmid so as to clone them by homologous recombination in yeast after co-transformation. The p*GPD*-*HDHD1*-DDAA vector was obtained by PCR amplification (oSL1155/1171) of the *HDHD1* DNA sequence obtained by gene synthesis using a specific 5’ primer carrying two D>A mutations and insertion of this insert into the pRS426 vector digested with EcoRI and BamHI as described above. This construct was verified by sequencing. The DDAA mutant was PCR amplified (oSL1297/oSL1298) and subcloned at NdeI/BamHI sites into pET15b-6His-HDHD1, a kind gift of Dr. E. Van Schaftingen, to substitute for endogenous HDHD1.

Plasmids expressing the wild-type *CYC8* gene and the mutant allele present in mutant #9 were obtained by PCR amplification on the corresponding genomic DNAs with primers (oSL1369/1370) containing a 50-bp overlap with a pRS415 vector (CEN, *LEU2*) digested with XbaI and BamHI, and co-transformation for cloning by homologous recombination in yeast as described above.

The plasmids generated in yeast were rescued by extraction (lithium acetate/SDS method (*104*)) and electroporation in bacteria, then amplified and sequenced before being re-transformed in the appropriate strains.

### Mass spectrometry and proteomics analyses

Samples used for the proteome-wide analysis of 2-deoxyglucose treatments were prepared from six liquid cultures (WT strain, BY4741) growing overnight at 30°C in 100mL of rich medium with 2% glucose to mid-log phase. On the next morning, 2-deoxyglucose was added to three of the cultures to a final concentration of 0.2% in order to obtain triplicates treated with 2-DG and triplicates without drug treatment (negative control). After 2.5 hours of incubation at 30°C, the six cultures were centrifuged at 4000*g* for 5 minutes at 4°C, resuspended in 500µL of 10% trichloroacetic acid (TCA, Sigma-Aldrich) and lysed by shaking after addition of glass beads (0.4-0.6mm, Sartorius) for 10 minutes at 4°C. Cell lysates were retrieved by piercing under the 1.5mL tubes and brief centrifugation. Precipitated proteins were centrifuged at 16000*g* for 10 minutes at 4°C, supernatants were discarded and pellets were rinsed 4 times in 1mL of 100% cold acetone.

Proteins were then digested overnight at 37°C in 20 μL of 25 mM NH_4_HCO_3_ containing sequencing-grade trypsin (12.5 μg/mL; Promega). The resulting peptides were sequentially extracted with 70% acetonitrile, 0.1% formic acid. Digested samples were acidified with 0.1% formic acid. All digests were analyzed by an Orbitrap Fusion equipped with an EASY-Spray nanoelectrospray ion source and coupled to an Easy nano-LC Proxeon 1000 system (all from Thermo Fisher Scientific, San Jose, CA). Chromatographic separation of peptides was performed with the following parameters: Acclaim PepMap100 C18 pre-column (2 cm, 75 μm i.d., 3 μm, 100 Å), Pepmap-RSLC Proxeon C18 column (50 cm, 75 μm i.d., 2 μm, 100 Å), 300 nl/min flow, using a gradient rising from 95 % solvent A (water, 0.1 % formic acid) to 40 % B (80 % acetonitrile, 0.1% formic acid) in 120 minutes, followed by a column regeneration of 20 min, for a total run of 140 min. Peptides were analyzed in the orbitrap in full-ion scan mode at a resolution of 120,000 (at *m/z* 200) and with a mass range of *m/z* 350-1550, and an AGC target of 2×10^5^. Fragments were obtained by higher-energy C-trap dissociation (HCD) activation with a collisional energy of 30 %, and a quadrupole isolation window of 1.6 Da. MS/MS data were acquired in the linear ion trap in a data-dependent mode, in top-speed mode with a total cycle of 3 seconds, with a dynamic exclusion of 50 seconds and an exclusion duration of 60 seconds. The maximum ion accumulation times were set to 250 ms for MS acquisition and 30 ms for MS/MS acquisition in parallelization mode.

Raw mass spectrometry data from the Thermo Fisher Orbitrap Fusion were analyzed using the MaxQuant software (*107*) version 1.5.0.7, which includes the Andromeda peptide search engine (*108*). Theoretical peptides were created using the *Saccharomyces cerevisiae* S288C proteome database obtained from Uniprot. Identified spectra were matched to peptides with a main search peptide tolerance of 6ppm. After filtering of contaminants and reverse identifications, the total amount of yeast proteins identified among the six samples was equal to 3425. Protein quantifications were performed using MaxLFQ (*109*) on proteins identified with a minimum amount of two peptides with a False Discovery Rate threshold of 0.05. LFQ values were then analyzed using Perseus (version 1.5.0.15). For the statistical analysis of yeast proteomes treated with 2-dexogylucose compared to negative control samples, each group of triplicates was gathered into a statistical group in order to perform a Student’s t-test. Results are presented in the form of Volcano-plots (*110*) and significant up-regulated and down-regulated candidates were determined by setting an FDR of 0.01 and an S_0_ of 2.

### GO-term analyses of proteomics data

The 79 significantly up-regulated candidates obtained in the proteomics analysis were used as input for the FunSpec web interface (http://funspec.med.utoronto.ca/) with default settings and a p-value cutoff of 0.01 in order to determine the Gene Ontology biological processes that are enriched in this list of 79 genes compared to the total *Saccharomyces cerevisiae* genome annotation (number of total categories=2062). The complete list of enriched GO biological processes with a p-value<0.01, as well as the genes included in each category, are displayed in Figure 1B.

### Protein extracts and immunoblotting

Yeast cultures used for protein extracts were all grown in synthetic complete medium. For each protein sample, 1.4mL of culture was incubated with 100µL of 100% TCA for 10 minutes on ice to precipitate proteins, centrifuged at 16000*g* at 4°C for 10 minutes and broken for 10 minutes with glass beads, as described for LC-MS/MS sample preparation. Lysates were transferred to another 1.5mL tube and centrifuged 5 minutes at 16000*g* at 4°C, supernatants were discarded and protein pellets were resuspended in 50µL*(OD_600_ of the culture) of sample buffer (50 mM Tris-HCl pH6.8, 100 mM DTT, 2% SDS, 0.1% bromophenol blue, 10% glycerol, complemented with 50mM Tris-Base pH8.8). Protein samples were heated at 95°C for 5 minutes and 10µL were loaded on SDS-PAGE gels (4-20%Mini-PROTEAN TGX Stain-Free, BioRad). After electrophoresis, gels were blotted on nitrocellulose membranes for 60 minutes with a liquid transfer system (BioRad), membranes were blocked in 2% milk for 20 minutes and incubated for at least two hours with the corresponding primary antibodies. Primary and secondary antibodies used in this study as well as their dilutions are listed in **Supplemental Table S5**. Membranes were washed three times for 10 minutes in Tris-Borate-SDS-Tween20 0.5% buffer and incubated for at least an hour with the corresponding secondary antibody (coupled with Horse Radish Peroxidase). Luminescence signals were acquired with the LAS-4000 imaging system (Fujifilm). Rsp5 was used as a loading control; alternatively, total proteins were visualized in gels using a trihalo compound incorporated in SDS–PAGE gels (stain-free TGX gels, 4–20%; Bio-Rad) after 1 min UV-induced photoactivation and imaging using a Gel Doc EZ Imager (Bio-Rad).

### Beta-galactosidase assays

β-Galactosidase assays were performed using 1mL of mid-log phase yeast cultures carrying the p_*DOG1*_-*LacZ* or p_*DOG2*_-*LacZ* plasmids, grown overnight in SC medium without uracil with glucose 2% and switched to the specified conditions. The OD (600 nm) of the culture was measured, and samples were taken and centrifuged at 16000*g* at 4°C for 10 minutes, cell pellets were snap frozen in liquid nitrogen and resuspended in 800µL of Buffer Z (pH=7, 50mM NaH_2_PO_4_, 45mM Na_2_HPO_4_, 10mM MgSO_4_, 10mM KCl and 38mM β-mercaptoethanol). After addition of 160µL of 4mg/mL ONPG (ortho-nitrophenyl-β-D-galactopyranoside, Sigma-Aldrich), samples were incubated at 37°C. Enzymatic reactions were stopped in the linear phase (60min incubation for p_*DOG2*_-*LacZ* and 120min incubation for the p_*DOG1*_-*LacZ* plasmid, as per initial tests) by addition of 400µL of Na_2_CO_3_, and cell debris were discarded by centrifugation at 16000*g*. The absorbance of clarified samples was measured with a spectrophotometer set at 420nm. β-Galactosidase activities (arbitrary units, AU) were calculated using the formula 1000*[A_420_/(A_600_* t)], where A_420_ refers to the enzyme activity and A600 is the turbidity of the culture, and t the incubation time. Each enzymatic assay was repeated independently at least three times.

### Drop tests

Yeast cells grown in liquid rich or synthetic complete medium for at least 6 hours at 30°C were adjusted to an optical density (600nm) of 0.5. Five serial 10-fold dilutions were prepared in 96-well plates and, using a pin replicator, drops were spotted on plates containing rich or SC medium containing 2% (w/v) agar and when indicated, 2DG (0.05%, or 0.2%, w/vol), sodium selenite (200µM) or tunicamycin (1µg/mL). Plates containing mannose (Fig 2H and 3G) were prepared as regular SC plates into which 2% mannose (w/vol) was added. Plates were incubated at 30°C for 3 to 4 days before scanning. For the adhesion test (Figure 6C), the plates were scanned and the colonies were then washed with tap water under a constant flow of water for 1 min as previously described (*75*). Excess water was removed before the plates were scanned again.

### 2-NBDG treatment and Fluorescence microscopy

For 2-NBDG treatments, yeast cells were grown overnight in SC medium containing 2% glucose to the exponential phase, then switched to SC medium with 0.5% lactate and incubated at 30°C for 30 minutes. This allowed the clearance of glucose from the medium, without which 2NBDG import was too weak to be observed since Glucose and 2NBDG compete for transport (*111*). 2NBDG (2-(*N*-(7-Nitrobenz-2-oxa-1,3-diazol-4-yl)Amino)-2-Deoxyglucose; Cayman Chemical) was then added at a concentration of 300µM for the indicated times.

Yeast were then rinsed twice in 1mL of cold water before being mounted on a glass slide and imaged at 25°C with a BX-61 fluorescence microscope (Olympus) equipped with a PlanApo 1.40NA 100X objective (Olympus), a QiClick monochrome camera (QImaging) and acquired using the MetaVue software (Molecular Devices). The 2-NBDG signal was visualized using a GFP filter set (41020 from Chroma Technology, Bellows Falls, VT; excitation HQ480/20X, dichroic Q505LP, emission HQ535/50m). Loa1-mCherry was visualized using an HcRedI filter set (41043 from Chroma Technology, Bellows Falls, VT; excitation HQ575/50X, dichroic Q610lp, emission HQ640/50m). Images were opened with ImageJ and processed for cropping and equivalent contrast adjustment.

### Isolation of spontaneous mutants and characterization

WT cells transformed with a p*DOG2*-LacZ plasmid (pSL410) were grown overnight in SC-Ura medium and ca. 2×10(6) cells were spread on SC-Ura plates containing 0.2% 2DG and grown for 6 days at 30°C. The clones obtained (24 clones) were restreaked on SC-Ura to isolate single clones. Resistance to 2DG was confirmed by drop tests (see Figure 5B). Beta-galactosidase enzyme assays and total protein extracts were performed on cultures grown to the exponential phase. For beta-galactosidase assays, the results of three independent experiments are shown along and were statistically tested using an unpaired t-test with equal variance.

Diploids were obtained by crossing each resistant mutant with a WT or *hxk2∆* strain of the opposite mating type (BY4742, Matα) and selecting single diploid clones on selective medium (SC-Met-Lys). Sequencing of the *REG1* and *HXK2* loci were done after PCR amplification on genomic DNA isolated from the corresponding clones. For whole genome sequencing, genomic DNA of the WT, clone 9 and clone 10 was purified using the Qiagen genomic DNA kit (Genomic-tip 20/G) using 30 OD equivalents of material following the manufacturer’s instructions after zymolyase treatment (Seikagaku). A PCR-free library was generated from 10 µg of gDNA and sequenced at the Beijing Genomics Institute (Hong Kong) on Illumina HiSeq 4000. The mutations in each clone were identified through comparative analysis of the variants detected by mapping their reads to the reference genome (BY4741) (*112, 113*) and those detected by mapping the WT reads to the reference. The differential variants were filtered by quality (vcf QUAL>1000) and manually inspected through IGV for validation (*114*).

### Cell culture and transfection

HeLa cells were maintained at 37°C and 5% CO_2_ in a humidified incubator and grown in Dulbecco’s modified Eagle’s medium (DMEM), supplemented with 10% fetal calf serum (FCS). Cells were regularly split using Trypsin-EDTA to maintain exponential growth. HeLa cells were transfected with plasmid pCMV-Sport6-HDHD1 and pCS2 (empty control) using Lipofectamine 2000 according to the manufacturer’s instructions. All culture media reagents were from Thermo Fisher Scientific. 2DG (Sigma) was used at a final concentration of 5 mM and tunicamycin (from *Streptomyces sp;* Sigma) was used at a final concentration of 5 µg/mL. For 2DG resistance assays, cells were grown in 10 cm2 flask and split in a 24-well plate in the absence or presence of 5 mM 2DG. Cells were counted each day with a hemocytometer after trypsinization and labeling with Trypan Blue. Total extracts were prepared by incubating cells (10 cm2) on ice for 20 min with 400 µL TSE Triton buffer (50 mM Tris-HCl pH 8.0, NaCl 150 mM, 0.5 mM EDTA, 1% Triton X-100) containing protease inhibitors (cOmplete protease inhibitor cocktail, EDTA-Free, Roche Diagnostics). Cells were then lysed mechanically with scrapers, and the lysate was centrifuged at 13,000 g, 4°C for 30 min. Proteins were assayed in the supernatant using the Bio-Rad Protein Assay reagent (Bio-Rad) and 40 µg proteins were loaded on SDS-PAGE gels.

### Recombinant His-tagged HDHD1 and HDHD1-DDAA protein purifications

*E. coli* BL21 bacteria were transformed with plasmids allowing the expression of His-tagged HDHD1 or HDHD1-DDAA. A 100mL-preculture was grown overnight in LB+Ampicillin (100 µg/mL), diluted 50-fold into 1L culture. The OD reached 0.7-0.9. IPTG (1 mM) was then added to induce the recombinant protein and cells were further grown at 25°C for 3 hours. Cells were harvested, the pellet was frozen in liquid N_2_ and thawed on ice. The pellet was resuspended in 20 mL of lysis buffer (HEPES 25mM pH 6.7, 300mM NaCl, imidazole 15 mM, beta-mercaptoethanol 2mM, glycerol 10% v/v and protease inhibitor cocktail [cOmplete protease inhibitor cocktail, EDTA-Free, Roche Diagnostics]). Cells were then sonicated and Triton X-100 was added to a final concentration of 1%. The lysate was centrifuged at 12,000rpm in a SW-32 rotor (Beckman Coulter) for 15min at 4°C, and then the supernatant was further centrifuged at 35 000 rpm for 1h at 4°C. The supernatant was incubated with 800 µL of Ni-NTA beads slurry (Qiagen) and rotated overnight at 4°C. The beads were collected by centrifugation (1000*g*, 2min, 4°C), resuspended in lysis buffer, and washed with 50 mL of lysis buffer at 4°C, and then washed again with 50 mL thrombin cleavage buffer (Hepes 50 mM, CaCl_2_ 5 mM, NaCl 100mM, glycerol 10%) at 4°C. The His-tag was removed by cleavage with 16 U of thrombin (Ref 27-0846-01, Sigma) added directly onto the beads for 2h at 25°C. The eluate was then collected and incubated with 500 µL benzamidin-sepharose 6B (GE Healthcare) at room temperature for 30 min to remove thrombin. The supernatant was collected and protein content was assayed by SDS-PAGE and colloidal blue staining (Brilliant Blue G-colloidal, Sigma), and protein concentration was assayed by the Bradford method (Bio-Rad protein assay, Bio-Rad).

### Enzyme assays

2-DG-6-Phosphate phosphatase assays were performed in 250 µL of reaction containing 1.5mM 2-DG-6-Phosphate (#17149, Cayman Chemicals, Ann Arbor, Michigan, USA) in 50 mM HEPES pH 6.7, 10 mM MgCl_2_ and 10% glycerol and 30 µg recombinant HDHD1 or HDHD1-DDAA. Samples were incubated at 37°C for various times (0, 5, 10, 15min) and the reaction was stopped by adding 150µL EDTA (0.5M). Then, the 2-deoxyglucose generated was assayed by adding 500µL of glucose assay reagent (GAGO20, Sigma) and further incubating at 37°C for 30min. The reaction was stopped by adding 500µL H_2_SO_4_ (12N) and the absorbance of the reaction was measured at 540 nm. A slope (A540 over time) was calculated to access to the enzyme activity and allowed to see that the reaction was in the linear range. The measurements were repeated 3 times.

### Statistical analysis

Mean values calculated using a minimum of three independent measurements from three biological replicates and are plotted with error bars representing standard error of the mean (SEM). Statistical significance was determined using a t-test for paired variables, as follows: *: P ≤ 0.05; **: P ≤0.01; ***: P ≤ 0.001; ns: P >0.05.

## Acknowledgments

We would like to thank Nicolas Joly for discussion about the HDHD1 *in vitro* assay, Gaelle Lelandais for help with Yeastract analysis, Pascual Sanz and Jose Antonio Prieto for the gift of the GST-Dog2 construct, Emile Van Schaftingen for the gift of the His-tagged HDHD1 construct, Peter Walter for the gift of the *pUPRE1*:lacZ reporter, David Levin for the gift of the pCYC(2xRlm1):lacZ reporter, Colin Stirling for the gift of the anti-invertase antibody and Martin Schmidt for advice regarding gDNA extraction for sequencing. We also thank Thibaut Léger and the Proteomics facility of the Institut Jacques Monod (supported by the Region Ile-de-France (SESAME), the Paris-Diderot University (ARS), and CNRS) for assistance. We thank Anna Babour, Alenka Čopič, Myriam Ruault, Eric Chevet, and members of the Léon and Chevet labs for insightful comments and critical reading.

## Author contributions

QD, AV, CL, SL contributed reagents, performed experiments, and acquired and analyzed the data; AF and JS analyzed the genome requencing data; SL and QD wrote the manuscript; SL directed the work.

## Data and materials availability

all data and material are available freely upon request.

## Supplemental material

Table S1. Proteomic response to 2DG treatment.

See Excel spreadsheet.

**Table S2.** Single nucleotide variants in clones #9 and #10 as compared to the WT strain, as identified by whole genome resequencing

| Clone | Chrom. | Position | Reference | Mutation | Gene | Mutation in coding region |
| --- | --- | --- | --- | --- | --- | --- |
| #9: | chrII | 464813 | G | A | *CYC8*<br>(*YBR112C*) | c.958C>T<br>(p.Gln320X) |
|  | chrXI | 570553 | C | G | *BET3*<br>(*YKR068C*) | c.357G>C<br>(p.Leu119Phe) |
|  | chrXII | 969100 | C | G | *YLR422W* | c.3204C>G<br>(p.Tyr1068X) |
| #10 | chrII | 464813 | G | A | *CYC8*<br>(*YBR112C*) | c.958C>T<br>(p.Gln320X) |
|  | chrIV | 283345 | T | A | Intergenic region *GET3-BUG1* (*YDL100C-YDL099W*) |  |
|  | chrVIII | 303712 | T | C | *TRA1*<br>(*YHR099W*) | c.952T>C<br>(p.Leu318Leu) |
|  | chrIX | 368278 | G | C | *PAN1*<br>(*YIR006C*) | c.1631C>G<br>(p.Thr544Ser) |
|  | chrXI | 570553 | C | G | *BET3*<br>(*YKR068C*) | c.357G>C<br>(p.Leu119Phe) |
|  | chrXIV | 381266 | A | C | *DGR1*<br>(*YNL130C-A*) | c.126T>G<br>(p.Cys42Trp) |
|  | chrXV | 499715 | G | A | *YOR093C* | c.2738C>T<br>(p.Ser913Leu) |

**Table S3.**
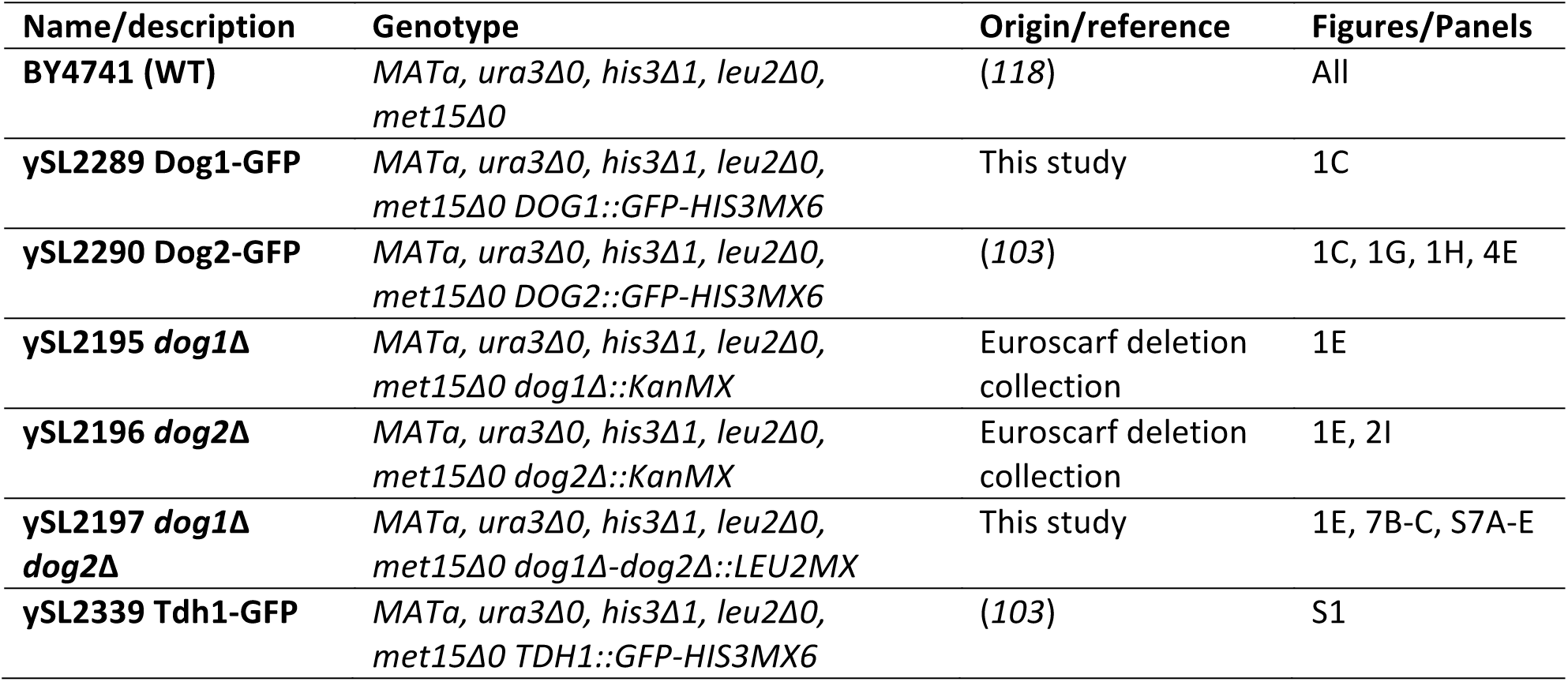
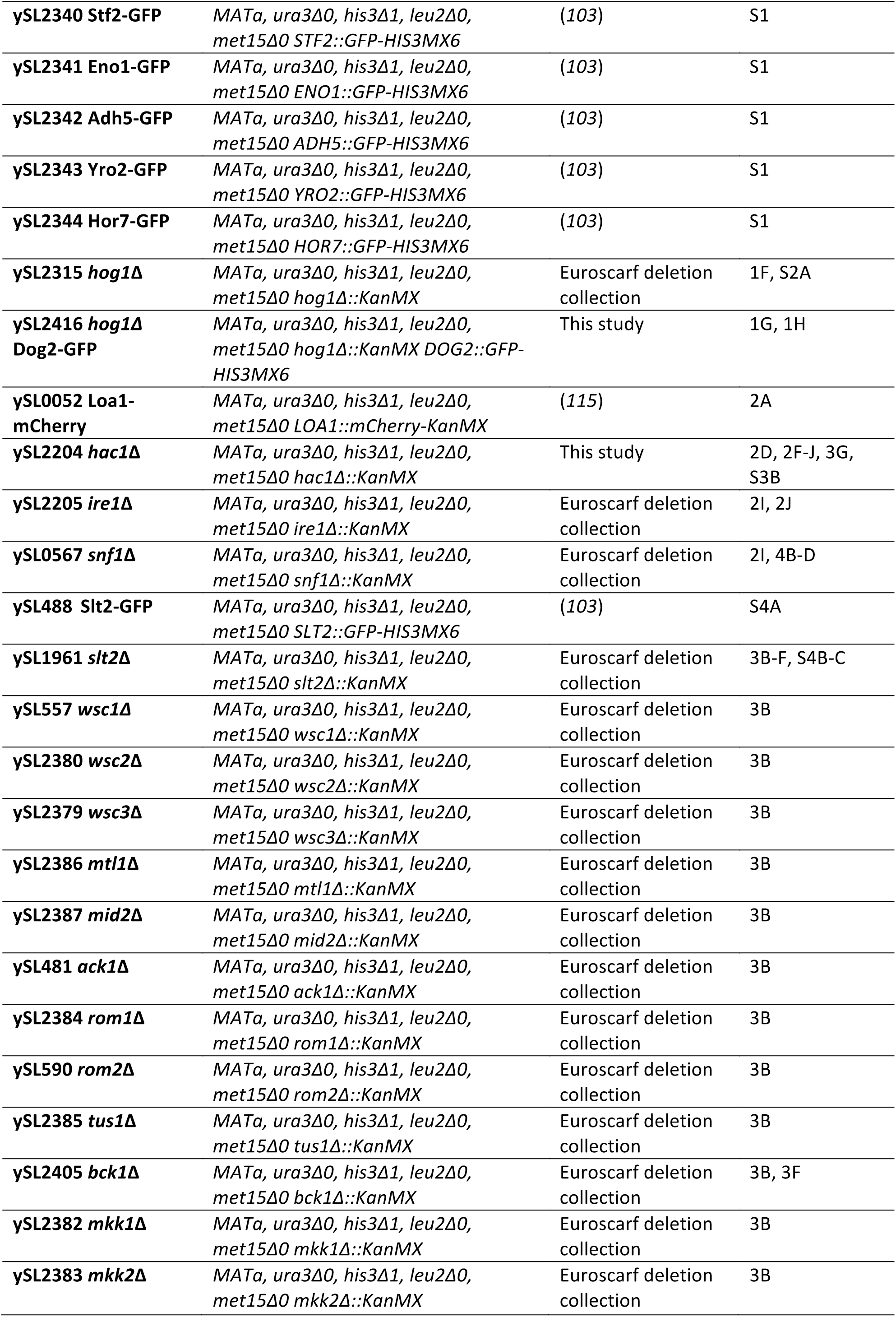
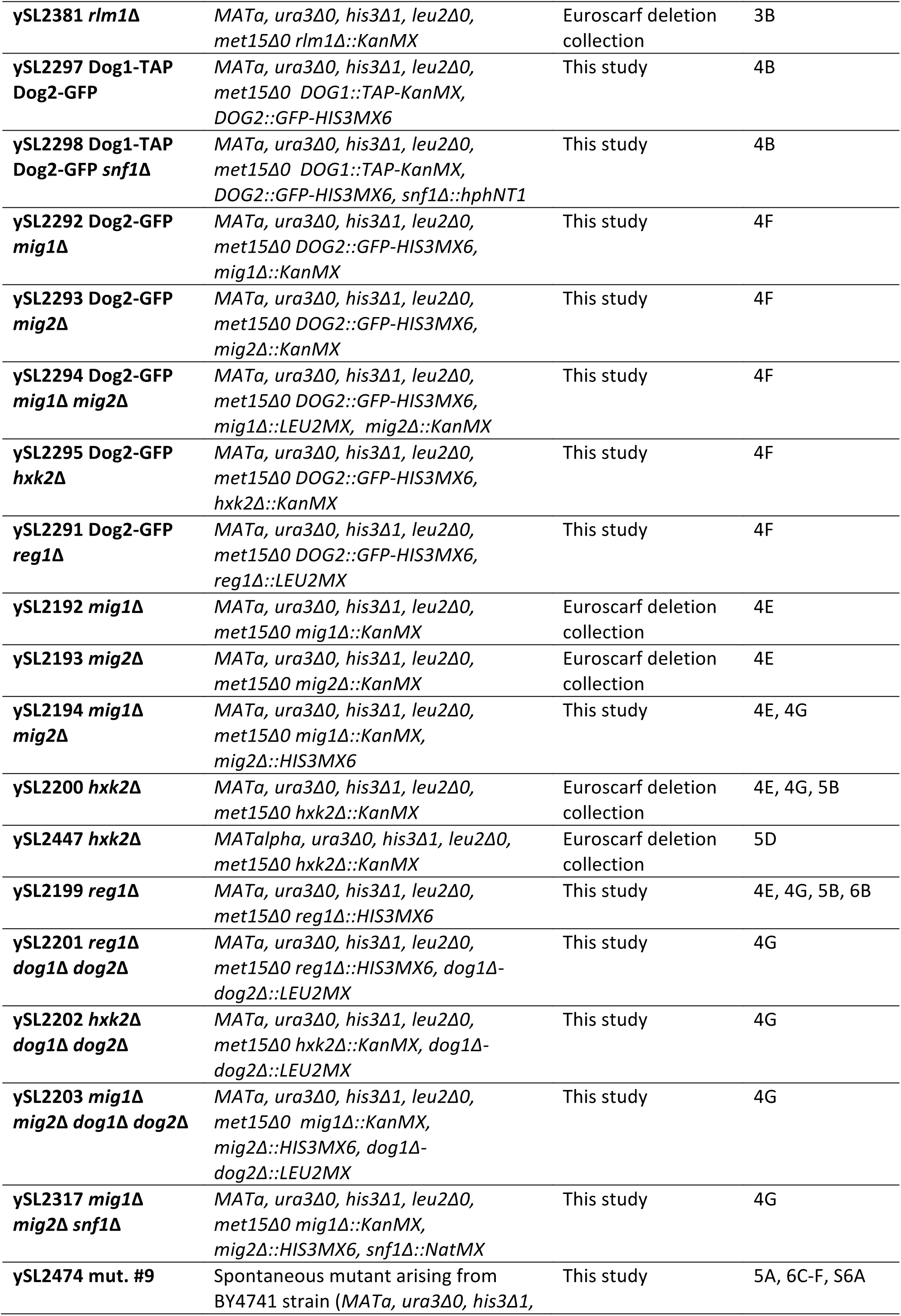
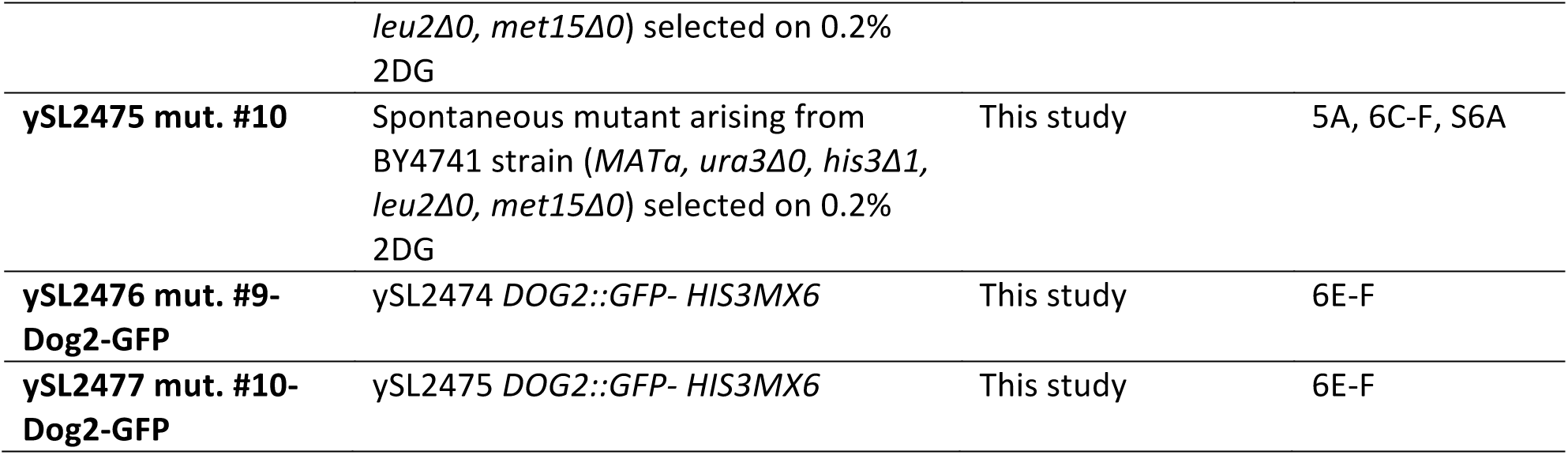
Yeast strains used in this study.

**Table S4.**
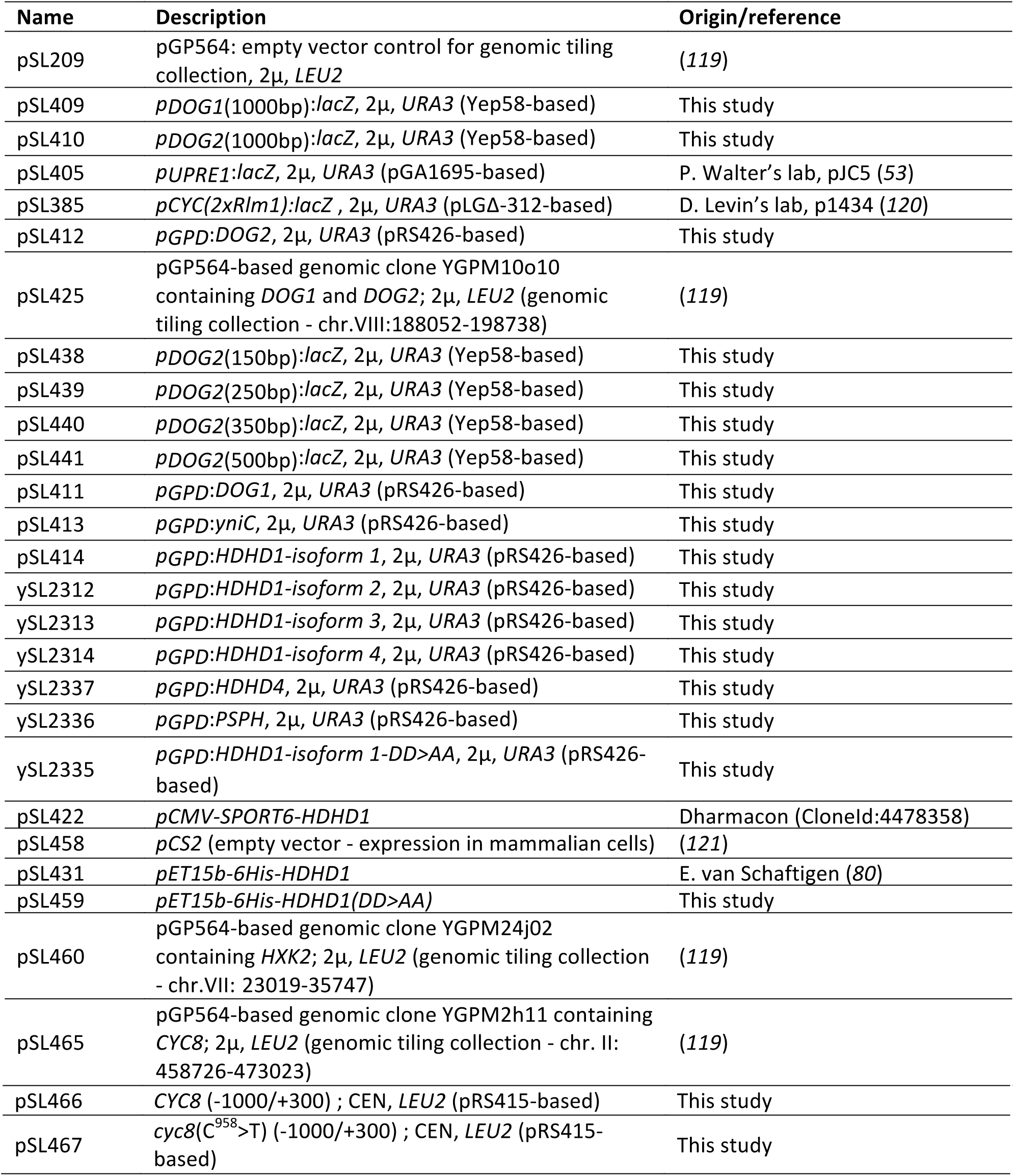
Plasmids used in this study.

**Table S5.**
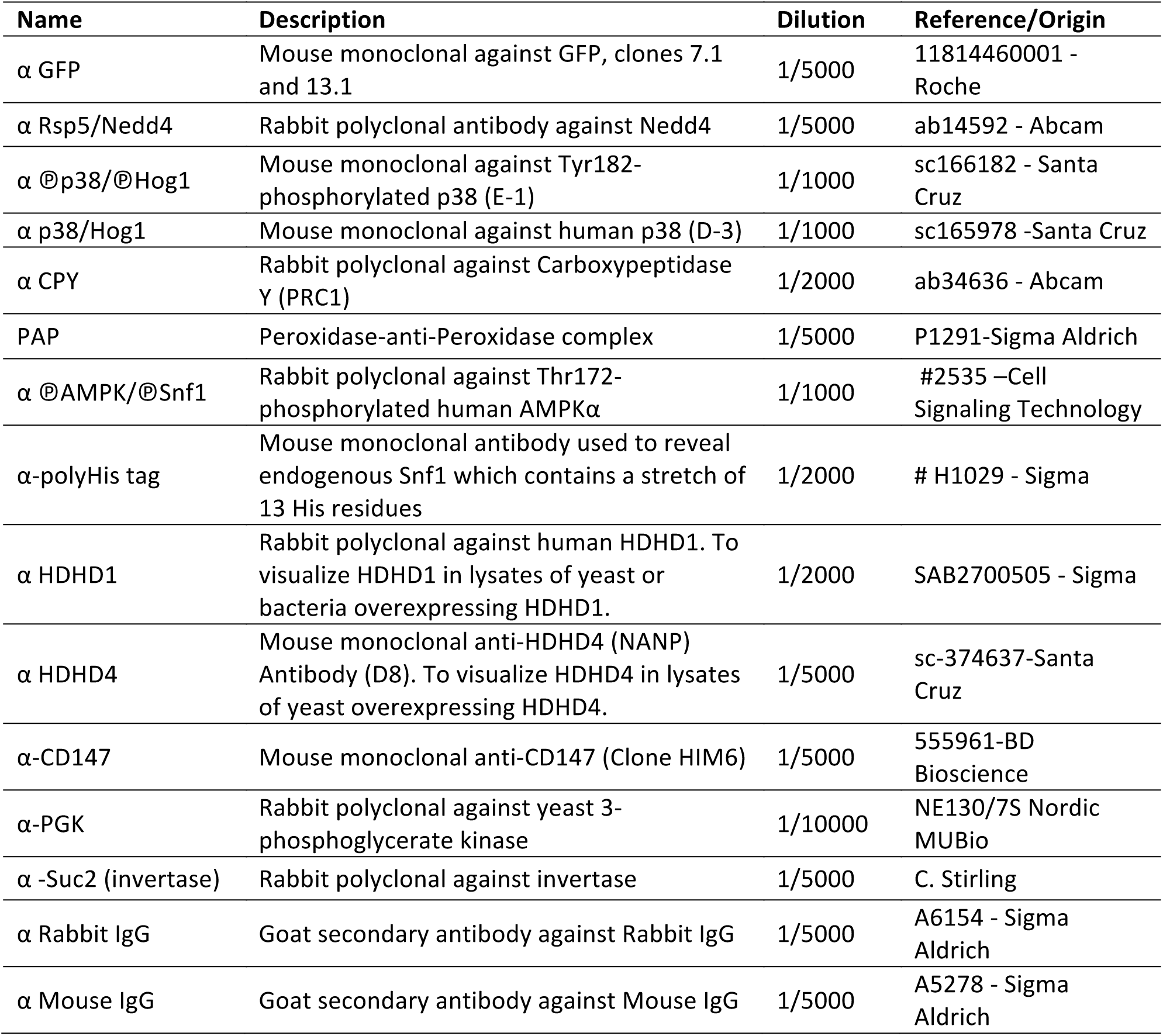
Antibodies used in this study.

**Figure S1.**
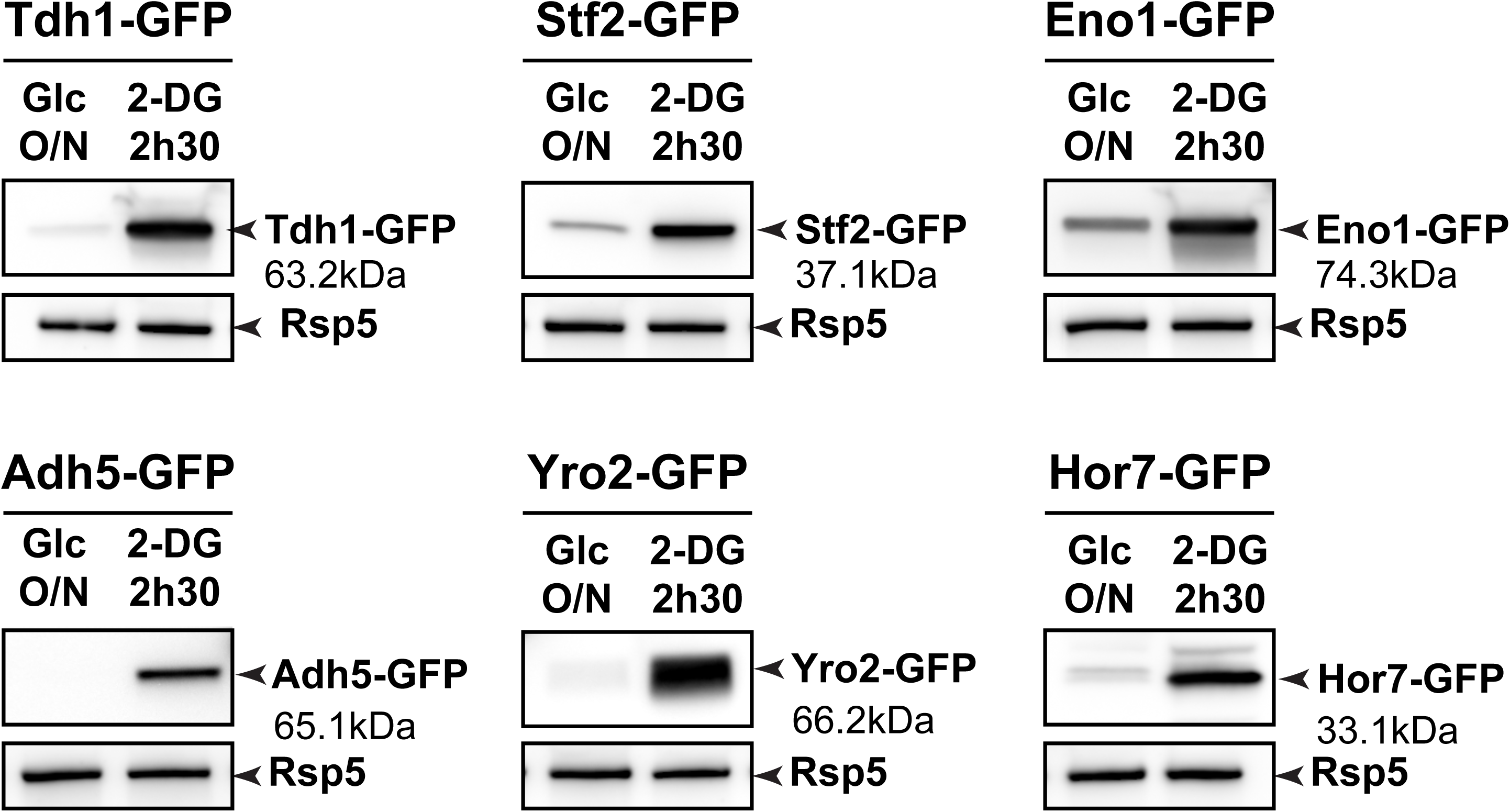
Characterization of other candidates upregulated by 2DG treatment. Western blot on total protein extracts of yeast cells expressing the indicated endogenously GFP-tagged proteins, before and after 2.5h 2DG addition, using an anti-GFP antibody. Rsp5 is used as a loading control.

**Figure S2.**
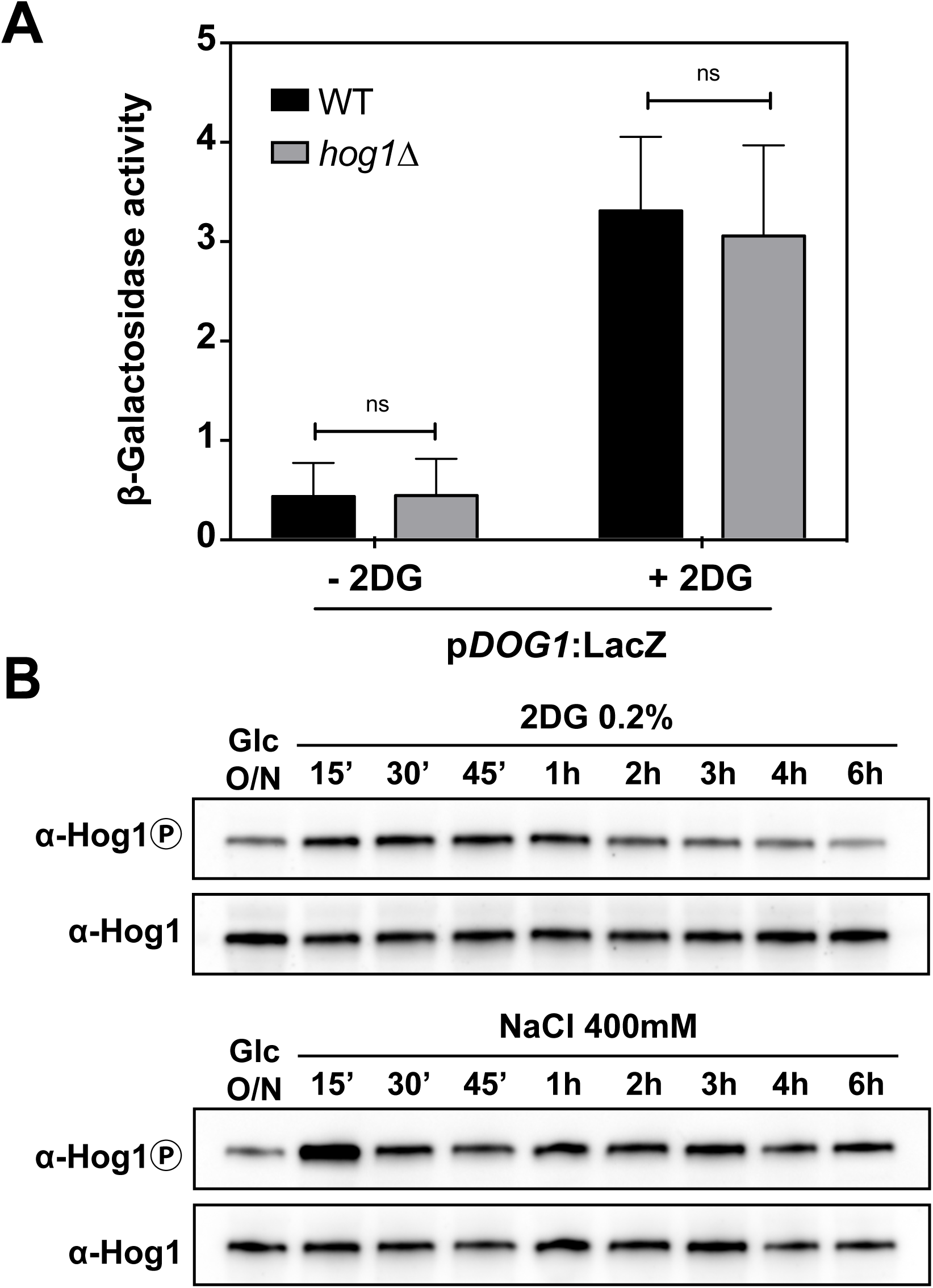
Hog1 signaling responds to 2DG but *DOG1* expression is not regulated by Hog1. (**A**). Beta-galactosidase assays of wild-type and *hog1∆* strains expressing *LacZ* under the control of the p*DOG1* promoter, before and after 2DG treatments for 3 hours. Error bars: SEM (*n*=3). (**B**). Western blot on total extracts of wild-type yeast cells grown overnight in SD medium and treated with 0.2% 2DG or 400mM NaCl for the indicated times. The anti-phospho-p38/Hog1 antibody enables the detection of phosphorylated Hog1 and the anti-p38/Hog1 antibody is used as a control for Hog1 abundance throughout the time course.

**Figure S3.**
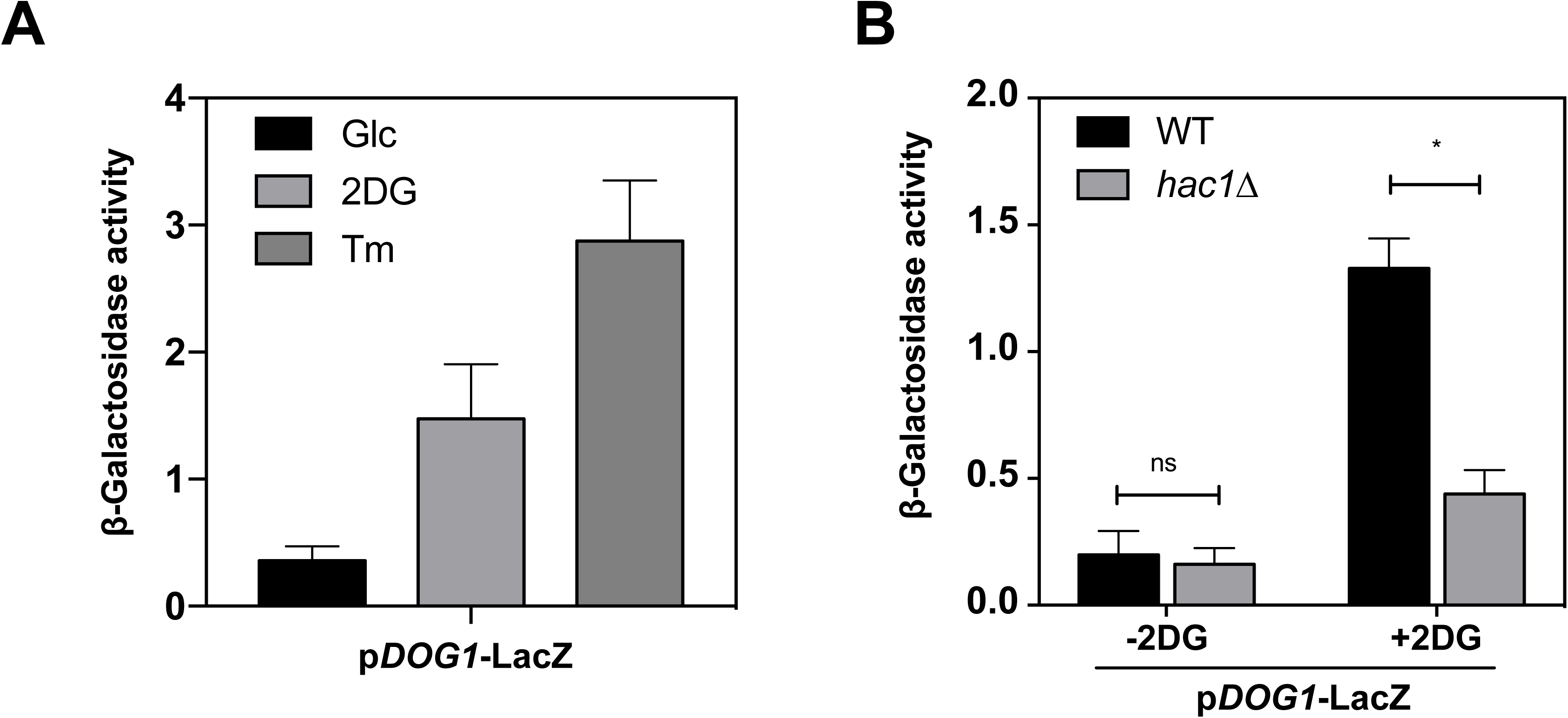
*DOG1* expression is regulated by the UPR pathway. (**A**). Beta-galactosidase assays on WT cells expressing *LacZ* under the control of the *DOG1* promoter and treated with 0.2% 2DG or 1µg/mL tunicamycin for 3 hours (± SEM, *n*=3). (**B**). Beta-galactosidase assays on WT and *hac1∆* cells expressing *LacZ* under the control the *DOG1* promoter, before and after 3h 2DG treatments (± SEM, *n*=3).

**Figure S4.**
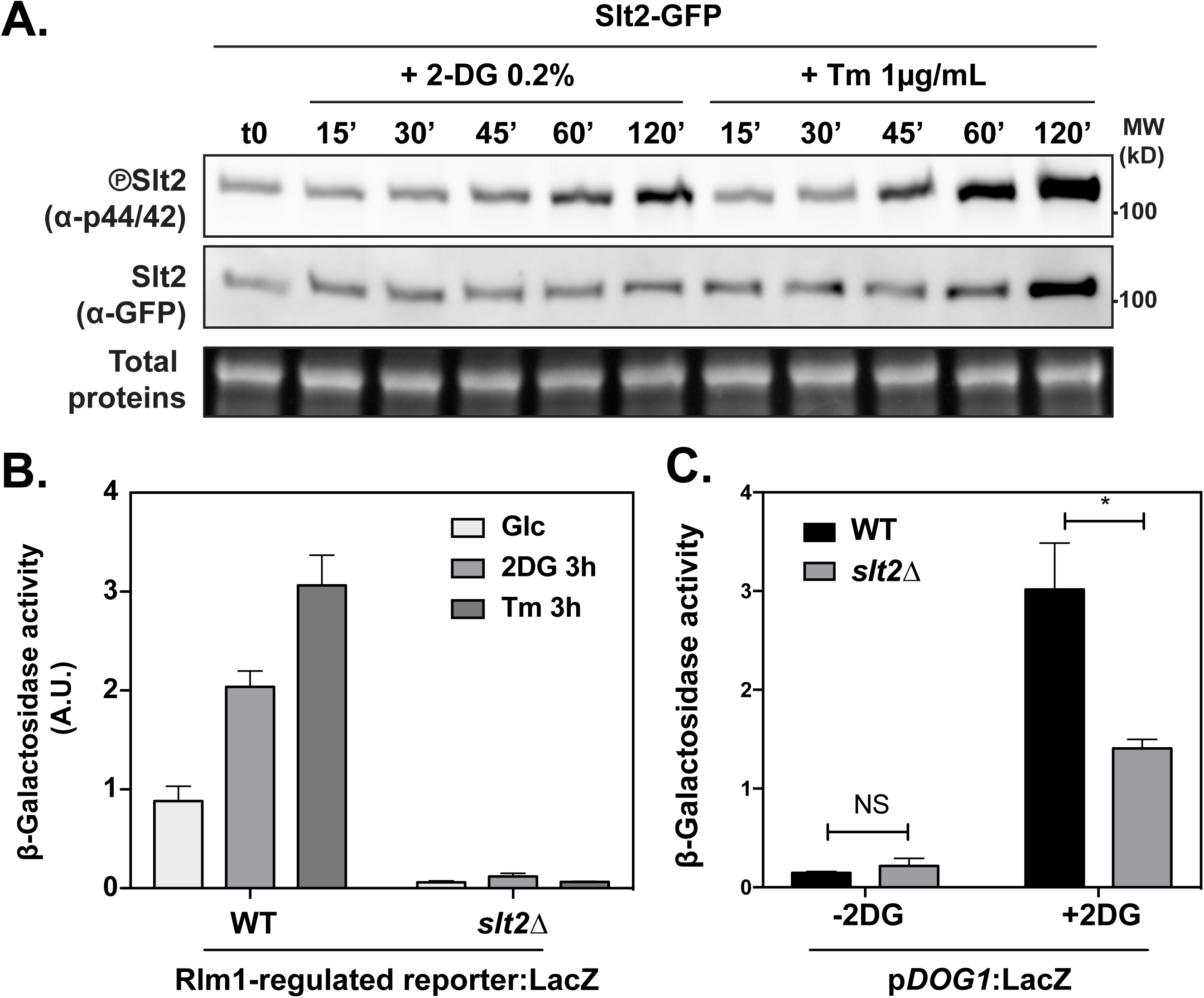
Slt2 participates in the regulation of *DOG1* expression. (**A**) A WT strain expressing an endogenously-tagged Slt2-GFP construct was grown in SC medium and treated with 2DG (0.2%) or tunicamycin (1 µg/mL). Total protein extracts were prepared at the indicated times and blotted with the following antibodies: anti-p44/42, to reveal activated (phosphorylated) Slt2, and anti-GFP to reveal total levels of Slt2. (**B**) Beta-galactosidase assays on WT and *slt2∆* cells expressing *LacZ* under the control of an Rlm1-regulated promoter, before and after 3h 2DG (0.2%) or tunicamycin (1 µg/mL) treatments (± SEM, *n*=3). (**C**) Beta-galactosidase assays on WT and *slt2∆* cells expressing *LacZ* under the control the *DOG1* promoter, before and after 3h 2DG treatments (± SEM, *n*=3).

**Figure S5.**
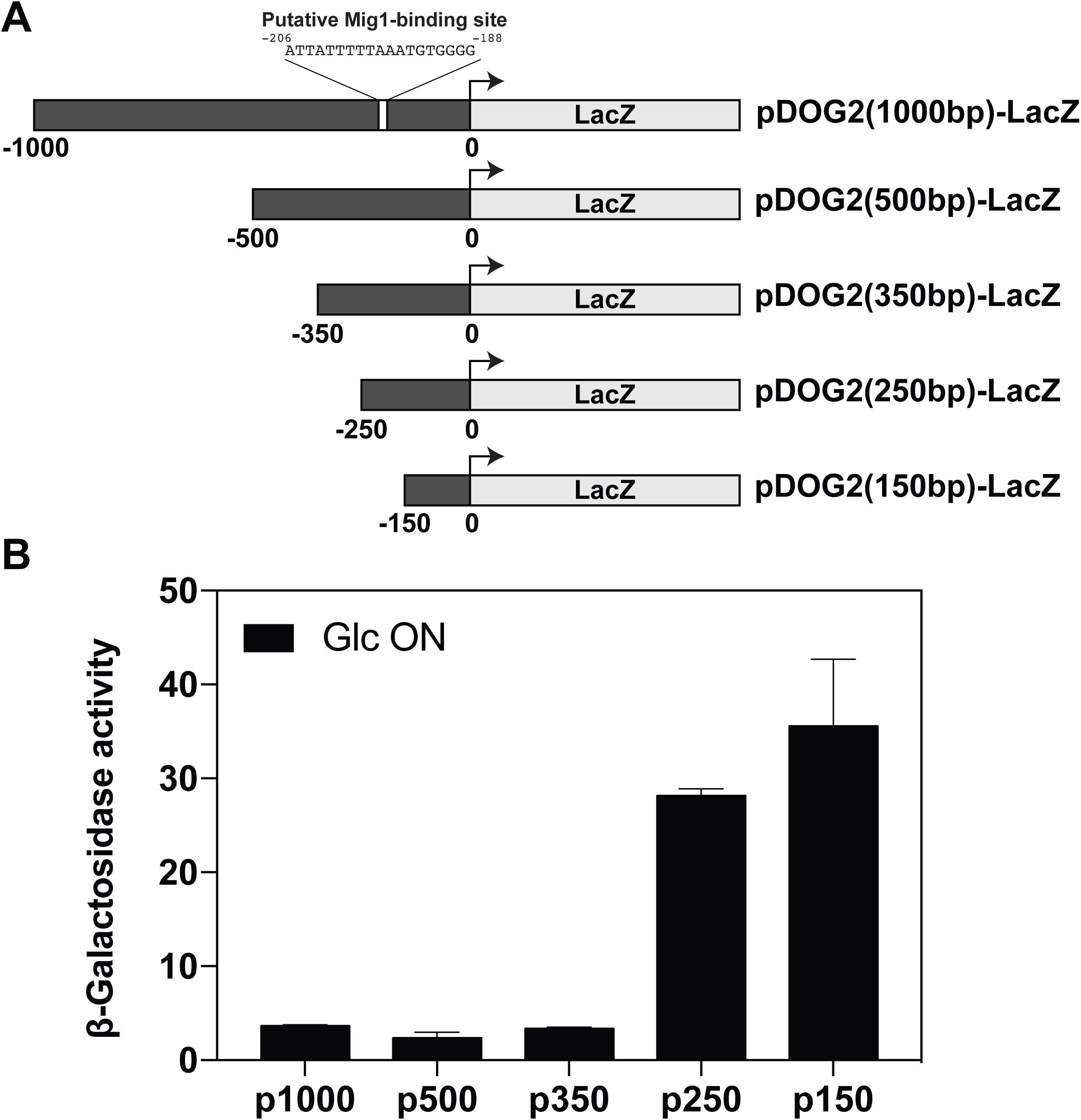
*Cis*-regions involved in the regulation of the *DOG2* promoter by glucose. (**A**). Schematic showing the various constructs generated to study *DOG2* promoter regulation by glucose availability. (**B**). Beta-galactosidase assays on WT cells expressing *LacZ* under the control of the indicated truncated version of the *DOG2* promoter after overnight growth in glucose medium (± SEM, *n*=3).

**Figure S6.**
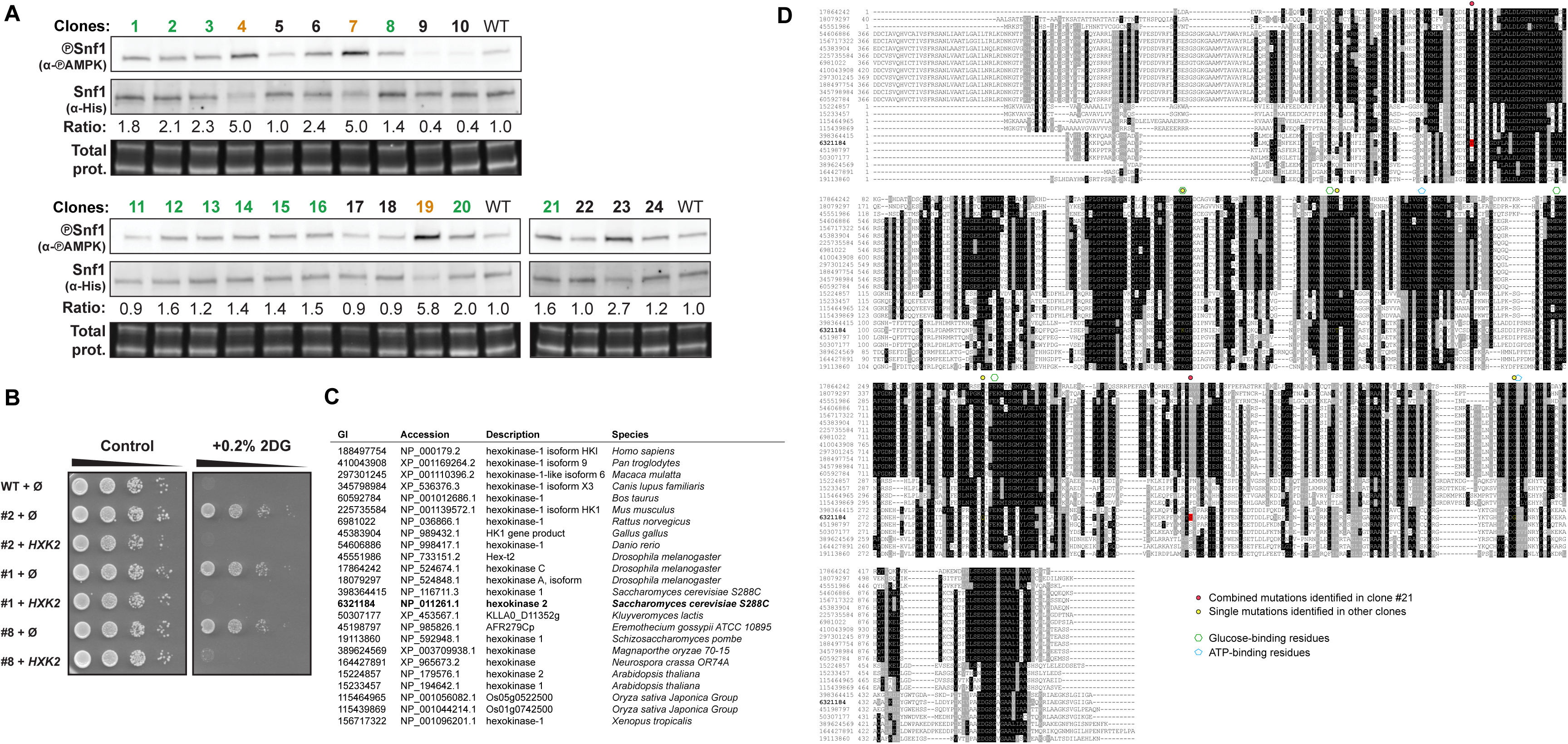
Multiple protein sequence alignment of Hxk2 orthologues and positions of the mutations identified. (**A**). A WT strain and the 24 resistant clones were grown in SC medium (exponential phase). Total protein extracts were prepared and blotted using antibodies allowing to reveal phosphorylated (activated) Snf1 (anti-phospho-AMPK) or total Snf1 (anti-polyHis tag, because Snf1 contains a stretch of 13 histidine residues that can be used for its detection) (*116*). The numbers below indicate the ratio of phospho-Snf1 signal over Snf1 signal, relative to the WT. (**B**). Growth test showing that the 2DG sensitivity of the 2DG-resistant clones #1 and #8 can be restored by expression of a multicopy, genomic clone containing *HXK2.* Clone #2 (which displays a mutation in *HXK2*, see Fig 5) is used as a positive control. (**C**). List of Hxk2 orthologues used for the alignment, as identified using HomoloGene entry #100530 (NCBI). (**D**). The sequences (numbers refer to their GI numbers) were aligned using ClustalX 2.1 (*122*) and formatted using BoxShade server (v.3.21). After the alignment, some sequences were truncated in Nt as indicated. The sequence in bold corresponds to that of *S. cerevisiae* Hxk2.

**Figure S7.**
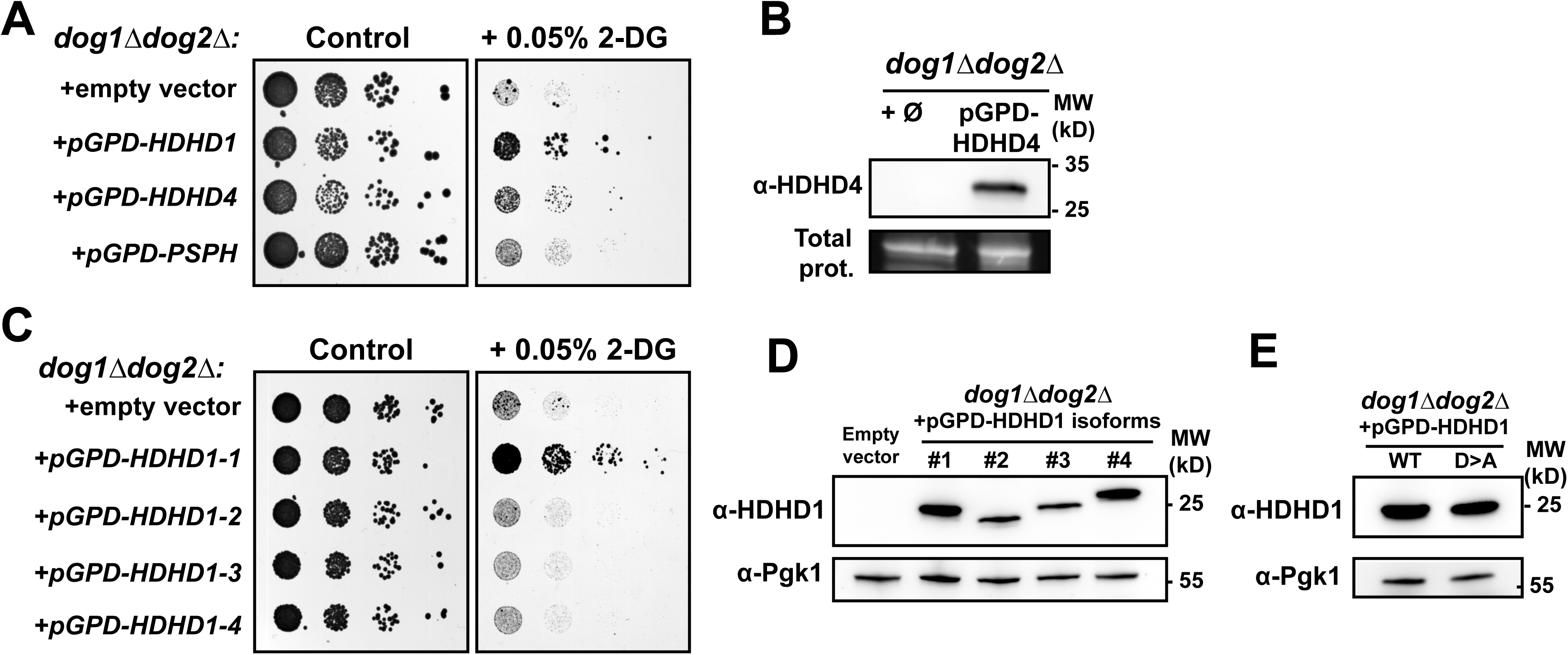
HDHD1 but not its close homologues HDHD4 nor PSPH allow resistance to 2DG. (**A**) Serial dilutions of *dog1∆dog2∆* strains transformed with the indicated plasmids were spotted on SC-Ura medium with or without 0.05% 2DG and were scanned after 3 days of growth at 30°C. (**B**) Total proteins extracts of *dog1∆dog2∆* cells transformed with an empty vector or a vector allowing the overexpression of HDHD4 were blotted with an anti-HDHD4 antibody. (**C**) Serial dilutions of *dog1∆dog2∆* strains transformed with an empty vector or the indicated vectors allowing the expression of the indicated HDHD1 isoforms were spotted on SC-Ura medium with or without 0.05% 2DG and were scanned after 3 days of growth at 30°C. (**D**) Total protein extracts were prepared from the same strains as in (**C**), and a western blot was revealed with an anti-HDHD1 antibody. The anti-Pgk1 antibody (phosphoglycerate kinase) was used as a loading control. (**E**) Total protein extracts were prepared from the same strains as in Fig 8C, and a western blot was revealed with an anti-HDHD1 antibody. The anti-Pgk1 antibody (phosphoglycerate kinase) was used as a loading control.

